# A toolbox for systematic discovery of stable and transient protein interactors in baker’s yeast

**DOI:** 10.1101/2022.04.27.489741

**Authors:** Emma J. Fenech, Nir Cohen, Meital Kupervaser, Zohar Gazi, Maya Schuldiner

## Abstract

Identification of both stable and transient interactions is essential for understanding protein function and regulation. While assessing stable interactions is more straightforward, capturing transient ones is challenging. In recent years, sophisticated tools have emerged to improve transient interactor discovery, with many harnessing the power of evolved biotin ligases for proximity labelling. However, biotinylation-based methods have lagged behind in the model eukaryote, *Saccharomyces cerevisiae,* possibly due to the presence of several abundant, endogenously biotinylated proteins. In this study, we optimised robust biotin-ligation methodologies in yeast and increased their sensitivity by creating a bespoke technique for downregulating endogenous biotinylation which we term ABOLISH (<u>A</u>uxin- induced <u>Bi</u>Otin <u>L</u>Igase dimini<u>SH</u>ing). We used the endoplasmic reticulum insertase complex (EMC) to demonstrate our approaches and uncover new substrates. To make these tools available for systematic probing of both stable and transient interactions, we generated five full-genome collections of strains in which every yeast protein is tagged with each of the tested biotinylation machineries; some on the background of the ABOLISH system. This comprehensive toolkit enables functional interactomics of the entire yeast proteome.

## Introduction

Cellular architecture and function require the action of protein machines often formed by protein complexes. Characterising the subunit identity of such complexes can be done using classic protein-protein interaction (PPI) assays such as immunoprecipitation (IP) of an epitope-tagged protein followed by mass spectrometry (MS) (Dunham *et al*, 2012). However, to uncover their protein substrates as well as their regulators (such as posttranslational modification enzymes) it is essential to probe transient interactions. Transient PPIs cannot easily be captured by such approaches since the nature of these experiments (such as the lysis conditions and long incubation times) select for only stable interactions between machinery subunits and cofactors. Therefore, when searching for transient PPIs between machineries and their clients or regulators, a more specialised approach is required.

One such approach is that of proximity labelling (PL) in which proteins proximal to a tested molecule are marked by a covalent tag that can be identified long after the interaction has ended. Biotin ligation represents a central PL method; with the first approach developed from the endogenous *Escherichia coli* biotin ligase, BirA (Cronan, 1990). BirA specifically biotinylates a lysine (K) residue within a short acceptor peptide sequence (Avi) (Beckett *et al*, 1999) in the presence of free biotin and ATP. Therefore, by tagging one protein with BirA and another with an Avi sequence (termed AviTag), stable and transient PPIs can be assessed in a pairwise manner using the high-affinity biotin binder, streptavidin, to detect biotinylated AviTag.

Having a pairwise assay enabled hypothesis-driven experiments but was less amenable to unbiased interactor discovery. Hence, a huge leap in the ability to utilise BirA-based methods for *de novo* discovery of interactions came with the creation of a promiscuous BirA mutant, BirA_R118G_ (Choi-Rhee *et al*, 2004). This mutant is able to biotinylate available K residues on accessible proteins without the requirement for a specialised acceptor sequence; making it possible to capture and identify multiple biotinylated interactors in one experiment using streptavidin affinity-purification (AP)-MS. Indeed, this powerful tool was shown to enable the discovery of new PPIs in mammalian cells (Roux *et al*, 2012). Later, a smaller version of BioID (BioID2) was generated from the *Aquifex aeolicus* BirA (Kim *et al*, 2016). However, the most active biotin ligase to date, TurboID, was produced by directed evolution of a BioID variant in *Saccharomyces cerevisiae* (from here on termed simply yeast) (Branon *et al*, 2018).

Surprisingly, despite the use of yeast to evolve TurboID, there has been limited use of these systems in yeast. BioID has so far only been applied to elucidate PPIs for ribosome- and mitoribosome-associated proteins (Opitz *et al*, 2017; Singh *et al*, 2020). One reason for this may be that BioID functions optimally at 37°C (Kim *et al*, 2016) and is minimally active at 30°C – the temperature at which yeast is grown. TurboID on the other hand, displays high activity at 30°C (Branon *et al*, 2018) and was employed to discover interactors for soluble cytosolic and exosomal proteins in the fission yeast, *Schizosaccharomyces pombe* (Larochelle *et al*, 2019); but remains untested for PPI discovery in *S.cerevisiae*. A more general reason explaining why biotin-based approaches have lagged behind in this powerful model organism is the presence of several highly expressed native proteins that are endogenously biotinylated (Sumrada & Cooper, 1982; Hasslacher *et al*, 1993; Brewster *et al*, 1994; Hoja *et al*, 2004; Kim *et al*, 2004; Nagaraj *et al*, 2012). These proteins hence make up a significant proportion of the signal following enrichment of biotinylated proteins and can therefore reduce the chance of identifying interactors – especially if they are of low abundance or transient.

Clearly, biotinylation based tools have immense power to uncover PPIs as has been demonstrated in multiple model systems, including mammalian cells (Roux *et al*, 2012; Go *et al*, 2021), mice (Uezu *et al*, 2016; Kim *et al*, 2021; Liu *et al*, 2021), flies (Uçkun *et al*, 2021), worms (Sanchez *et al*, 2021; Artan *et al*, 2021) and plants (Zhang *et al*, 2019; Mair *et al*, 2019). This widespread utilisation incentivised our work to make this tool applicable for systematic identification of PPIs, particularly transient ones, in yeast. To this end, we address the current gap in PL technology in yeast by optimizing protocols for discovery of stable and transient interactions using a variety of biotin ligation-dependent techniques. Moreover, we develop a novel approach, which we term ABOLISH (<u>A</u>uxin-induced <u>B</u>i<u>O</u>tin <u>LI</u>gase dimini<u>S</u>Hing), for downregulation of endogenous biotinylation to increase the signal-to-noise ratio and make PPI discovery more robust. We showcase the power of these approaches by uncovering a set of new substrates for the endoplasmic reticulum (ER) localised insertase, the ER membrane complex (EMC). Most importantly, to enable these powerful tools to be used easily and rapidly by the entire yeast community and to promote systematic probing of interactions, we generated a collection of full-genome libraries in which each yeast gene is preceded by either TurboID-HA, BioID2-HA, BirA or AviTag; with the ABOLISH system integrated into several of them. Altogether these freely available libraries provide a powerful platform for high-content PPI screening and ultimately substrate recognition and protein function discovery in yeast.

## Results

### Developing ABOLISH - a strategy to enhance the signal-to-noise ratio for exogenous biotin ligases

To expand the arsenal of biotin-based tools available for protein interaction profiling in yeast it is essential to take into consideration endogenously biotinylated proteins (Sumrada & Cooper, 1982; Hasslacher *et al*, 1993; Brewster *et al*, 1994; Hoja *et al*, 2004; Kim *et al*, 2004). This is because most biotinylated yeast proteins are highly abundant (Figure 1A) and therefore can mask many of the expected PPI assay signals on a streptavidin blot (Figure 1B) or take up a large percent of the reads from MS analyses. Since PL relies on biotinylated protein enrichment and detection, it would clearly be advantageous to reduce the background signal from the endogenously biotinylated proteins. To do this, we created a new method of endogenous biotinylation reduction that we call ABOLISH, for <u>A</u>uxin-induced <u>B</u>iOtin <u>LI</u>gase dimini<u>S</u>Hing. In this method, Bpl1, the only endogenous yeast biotin ligase, is C-terminally tagged with an auxin-inducible degron (AID*, (Nishimura *et al*, 2009; Morawska & Ulrich, 2013)). Therefore, in the presence of auxin and the *Oryza sativa* transport inhibitor response 1 (OsTIR1, (Nishimura *et al*, 2009)) adaptor protein, the controlled and transient degradation of this essential enzyme ensues, leading to a reduction in biotinylation of its substrates. Indeed, growth in reduced-biotin (RB) media (Jan *et al*, 2014), followed by auxin addition to induce Bpl1 degradation, and finally treatments with a biotin pulse (illustrated in Figure EV1A) demonstrated that while RB media dramatically reduced endogenous biotinylation levels, this was rapidly reversed upon addition of free biotin (Figure EV1B). However, this reversal was not observed if auxin was used to deplete Bpl1-AID*-9myc; proving that ABOLISH reduces background biotinylation noise.

**Figure 1:**
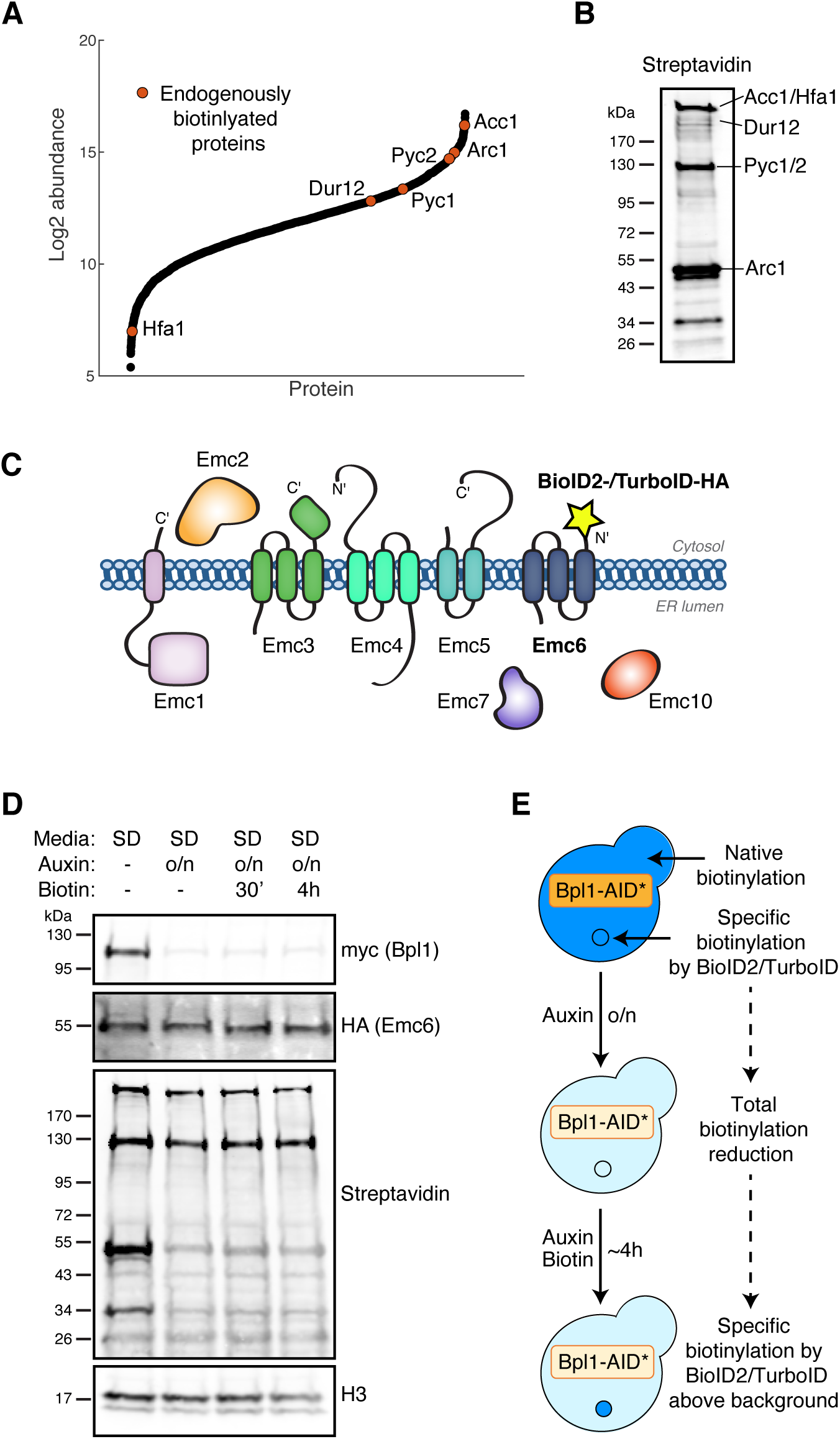
Utilising an auxin-inducible degron to reduce levels of endogenous biotinylation in TurboID-expressing cells. (A) The abundance of endogenously biotinylated proteins (orange circles) relative to the abundance of all yeast proteins (black circles) as determined by (Nagaraj *et al*, 2012). (B) A streptavidin blot of a control strain showing the running pattern of the endogenously biotinylated proteins highlighted in (A), and labelled according to (Pirner & Stolz, 2006). (C) Schematic of the EMC showing Emc6 N-terminally tagged with BioID2-HA or TurboID-HA. (D) Western blot analysis of a strain expressing TurboID-HA-Emc6, Bpl1- AID*-9myc and OsTIR1 which was grown overnight in SD media with or without auxin. The cells were then diluted into fresh SD media and grown to mid logarithmic phase (about 4hours) with or without auxin (respectively). When required, biotin was added either 30min before harvesting, or for the entire 4hours. H3 (histone H3) is used as a loading control. (E) Schematic of the growth conditions used in (D) which were selected for the remaining ABOLISH experiments.

Next, we wanted to understand how this system would interact with exogenous promiscuous biotin ligases. To do that, we chose a complex for which we could follow both stable and transient PPIs: the most recently characterised ER-resident insertase; the ER membrane protein complex, EMC (Guna *et al*, 2018). This highly conserved machinery is composed of eight subunits (Emc1-7 & Emc10) in yeast (Jonikas *et al*, 2009) and 10 (EMC1-10) in humans (Christianson *et al*, 2012). Since its discovery as an insertase for moderately hydrophobic tail-anchor (TA) proteins (Guna *et al*, 2018; Volkmar *et al*, 2019), it has also been found to insert multi-pass transmembrane domain (TMD)-containing proteins into the ER (Shurtleff *et al*, 2018; Chitwood *et al*, 2018; Tian *et al*, 2019; Bai *et al*, 2020). Furthermore, it is required for the biogenesis of single-pass TMD proteins which do not contain a signal peptide (also known as type III membrane proteins, (O’Keefe *et al*, 2021)) and transmembrane proteins which traffic from the ER to lipid droplets (LD) (Leznicki *et al*, 2021). To this end, it has a wide (and not yet fully characterised) substrate range and a clear set of stable interactions.

To test the capacity of biotin ligases to label both stable and transient interactions, and to evaluate whether ABOLISH enhances the detection of these labelled proteins, we tagged Emc6 at its N-terminus with either BioID2-HA or TurboID-HA (Figure 1C). A third strain expressing both TurboID-HA-Emc6 and the ABOLISH system was also generated, along with three control strains in which Sbh1 (an independent ER membrane protein), rather than Emc6, was tagged. All promiscuous biotin ligase tags were preceded by the constitutive moderate *CYC1* promoter, and all tagged proteins ran at their expected molecular weights as determined by SDS-PAGE (Figure EV1C).

Surprisingly, we found that overnight growth in RB media resulted in a decrease in the amount of TurboID-HA-tagged proteins (Figure EV1D); thus negating the signal-to-noise advantage conferred by ABOLISH. To uncover a condition where the TurboID-tagged protein levels are not reduced but endogenous biotinylation levels are, we tested several different parameters and found that overnight treatment with auxin in SD media (Figure EV1E, 2^nd^ lane) resulted in the strongest background biotinylation reduction without a loss in TurboID-HA-Emc6. Interestingly, the abundance of the ER translocon, Sec61, remained constant independent of the conditions. This suggests that the loss of TurboID-tagged proteins triggered by biotin depletion may be a regulatory adaptation to biotin starvation, rather than general protein degradation from the ER. Finally, to ensure sufficient labelling material and time for true PPI events to be captured by TurboID, biotin was added for 30 minutes or 4 hours prior to collecting the cells (Figure 1D). Importantly, even 4h of exogenous biotin addition did not negate the effect of the auxin-induced depletion of endogenous biotinylated proteins and therefore this pipeline was adopted for future ABOLISH experiments (illustrated in Figure 1E). These data collectively demonstrate that the ABOLISH method can be harnessed to reduce background biotinylation ‘noise’, paving the way for enhanced signal detection from exogenous proximity-labelling enzymes.

### Comparing three biotin ligase systems identifies their ability to uncover both stable and transient protein-protein interactions by LC-MS/MS

While IPs of epitope-tagged proteins enrich for stable interactors (in this case EMC complex components), streptavidin APs should capture transient interactions labelled by exogenous biotin ligases (in this case clients inserted into the ER by the EMC) as well as a subset of stable ones. To directly compare the type of interactions that we can identify we analysed, either by HA-IP or streptavidin-AP, strains expressing either BioID2-HA-Emc6, TurboID-HA- Emc6, or TurboID-HA-Emc6 on the background of the ABOLISH system (Figure 2A).

**Figure 2:**
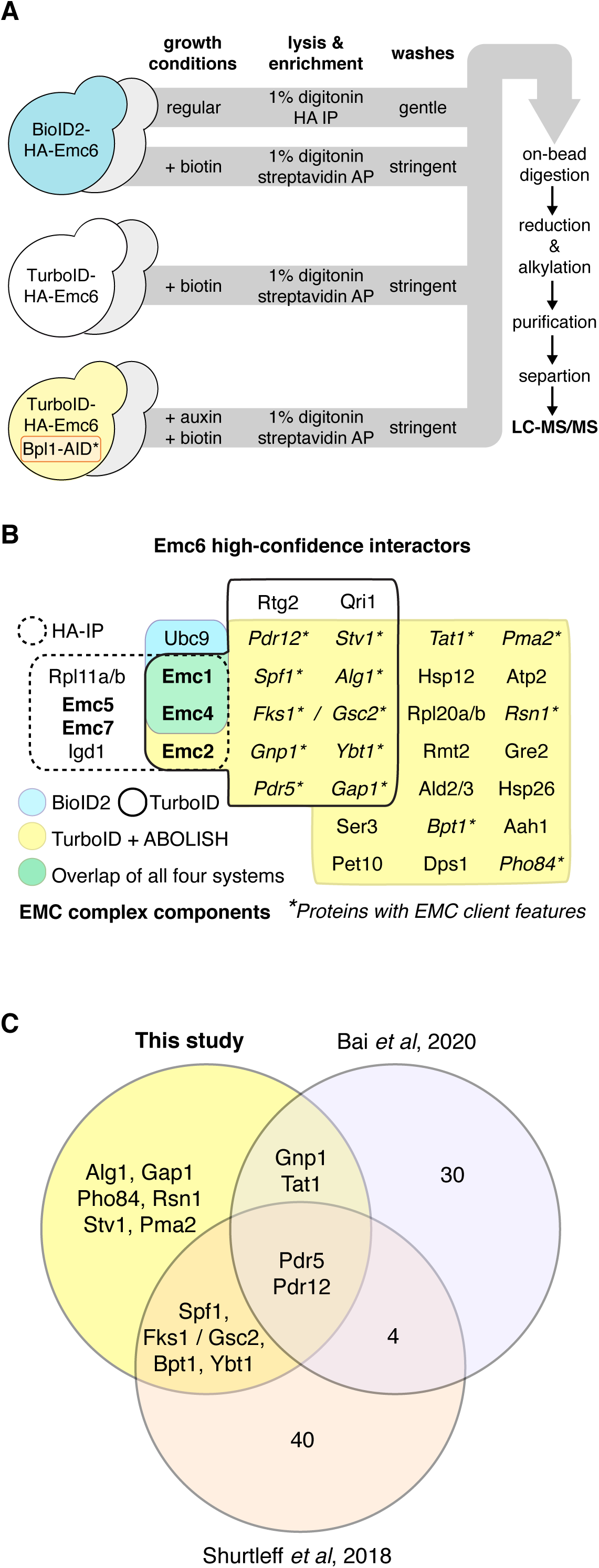
Finding stable and transient Emc6 interactors by a multi-faceted LC-MS/MS approach. (A) Schematic of the workflow used for LC-MS/MS sample preparation of the BioID2-HA-Emc6, TurboID-HA-Emc6, and TurboID-HA-Emc6/ABOLISH strains and their control counterparts (depicted as grey yeast). All strains were prepared in biological triplicate. (B) High-confidence interacting proteins of Emc6 determined by: HA-IP of BioID2-HA-Emc6 (dotted outline); streptavidin-AP of BioID2-HA-Emc6 (blue fill); streptavidin-AP of TurboID-HA-Emc6 (black, solid outline); and streptavidin-AP of TurboID-HA-Emc6/ABOLISH (yellow fill). EMC complex members and proteins with classical features of EMC substrates are marked in bold and with asterisks, respectively. (C) Overlap between the proteins highlighted with asterisks in (B) and yeast EMC clients found by two independent studies (Shurtleff *et al*, 2018; Bai *et al*, 2020).

All Emc6 samples were compared to their Sbh1 control counterparts to find high-confidence interacting proteins (Table S1 and via PRIDE, PXD033348). These were defined by the following criteria: a *p-value* of ≤0.05 (streptavidin samples) or ≤0.1 (HA samples); a fold-change of ≥2; and identification by two or more unique peptides. From the HA-IP, as expected, almost the entire EMC complex satisfied these requirements (Figure 2B; dotted outline). From the streptavidin-AP samples, only three high-confidence interactors were found by BioID2, two of which were Emc1 and Emc4 (Figure 2B; blue fill). This confirms that although BioID2 is able to label *bona fide* interactors, its capacity is limited likely due to its relatively low catalytic activity at 30°C (Kim *et al*, 2016). TurboID, on the other hand, identified 14 high-confidence interactions (Figure 2B; black, solid outline). Looking at stable interactors, Emc2 was found in addition to Emc1 and Emc4, already hinting at increased labelling functionality relative to BioID2. However, most encouragingly, of the remaining 11 putative interactors, nine had membrane protein features classically associated with EMC clients, suggesting an increased capacity to uncover transient interactions (Figure 2B, asterisks). Eight of these are multi-pass TMD secretory pathway proteins, and the remaining protein (Alg1) is a lipid-droplet (LD) protein with a single N-terminal TMD – a characteristic recently demonstrated to define EMC-dependence (Leznicki *et al*, 2021).

Furthermore, incorporating the ABOLISH system enabled detection of even more putative interactors labelled by TurboID (Figure 2B; yellow fill). The overlap between both TurboID strategies was very large with the same stable interactions and all nine candidate substrates being found. Another 16 high-confidence interaction partners were identified, five of which were secretory pathway multi-pass TMD proteins (Figure 2B; asterisks). Comparing our list of putative substrates to published yeast EMC client data found by ribosome-profiling (Shurtleff *et al*, 2018) and proteomic analysis of WT vs *EMC3* KO cells (Bai *et al*, 2020), revealed that eight out of the 14 identified had previously been found in either study (Figure 2C) supporting the validity of our transient PPI discovery. To our knowledge, this is the first time TurboID has been successfully used in baker’s yeast, and our data demonstrate that it labels both stable and transient PPIs. In addition, the ABOLISH system enhances the capacity to detect TurboID-labelled interactors.

### Validating new EMC substrates using genetic tools and a natively-expressed pairwise biotinylation method

The similarity between our list of candidate EMC clients and published datasets (Figure 2C) strongly suggested that TurboID-mediated proteomics, both with and without ABOLISH, identified *bona fide* EMC substrates. Such substrates should therefore be affected by loss of the complex and indeed it was previously shown that the abundance and/or localisation of true EMC substrates changes upon Emc3 loss (Bai *et al*, 2020). We therefore deleted *EMC3* on a selection of our candidates and observed that GFP-Pdr12 changed its localization relative to the control strain (Figure 3A, top panel). Pdr12 is a plasma membrane ATP-binding cassette (ABC) transporter which first requires insertion into the ER before trafficking to its final destination. Therefore, the accumulation of Pdr12 on the ER in the Δ*emc3* strain likely signifies a pre-inserted Pdr12 population at the ER surface. Deletion of *EMC3* also strongly reduced the abundance of GFP-Alg1 (Figure 3A, middle panel) and, to a lesser but still significant extent, Gnp1-GFP (Figure 3A, bottom panel; quantified in 3B). These functional assays support these proteins as newly validated clients of the EMC complex.

**Figure 3:**
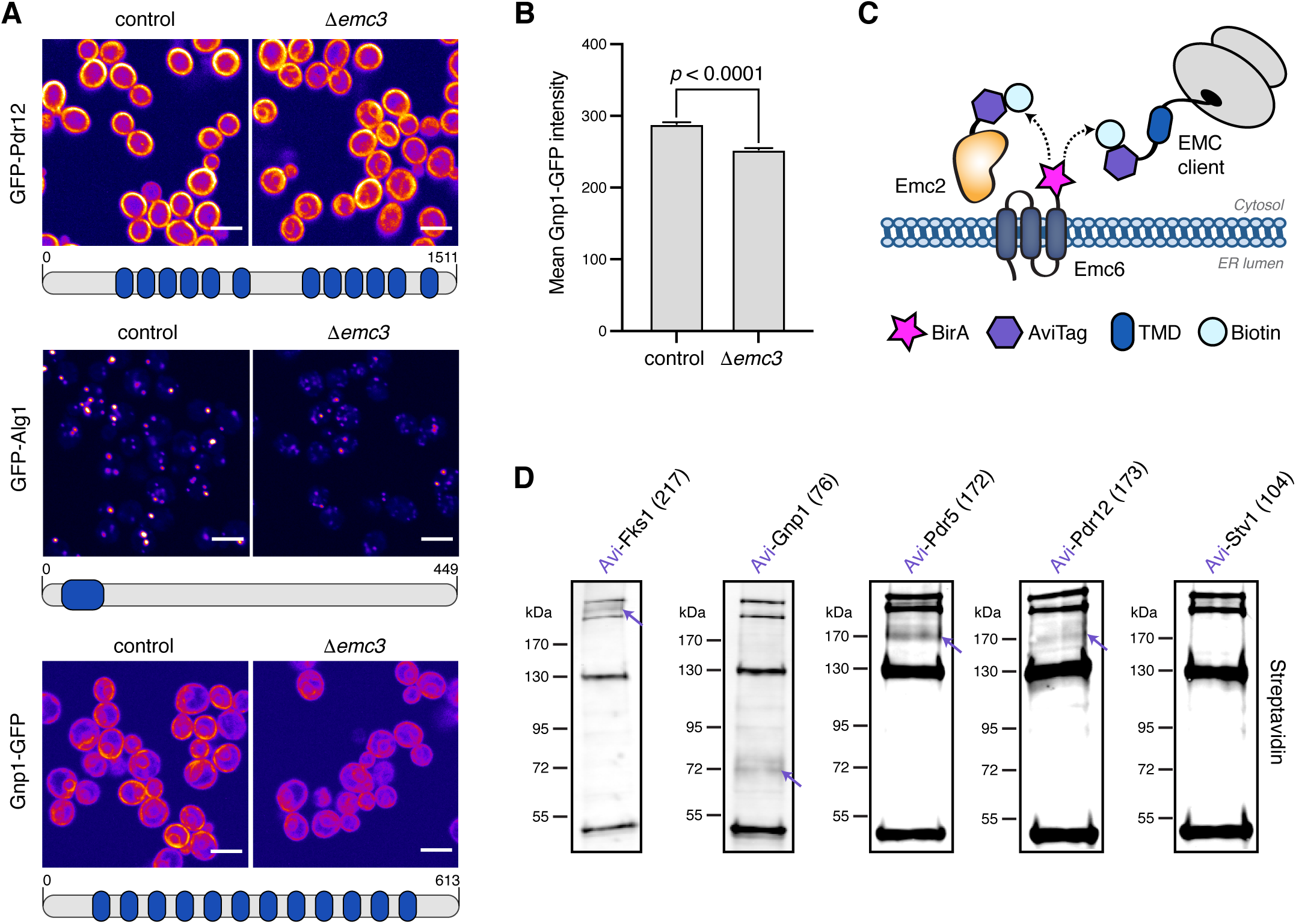
Verifying new EMC clients using microscopy and BirA-AviTag combined with ABOLISH. (A) Fluorescence microscopy images of control or Δ*emc3* strains containing either Pdr12 N’ tagged with GFP, Alg1 N’ tagged with GFP, or Gnp1 C’ tagged with GFP. Topological schematics of Pdr12, Alg1 and Gnp1 feature beneath their respective images, with TMDs highlighted in blue (Weill *et al*, 2019). All images are from two independent biological repeats and are digitally coloured using the ‘fire’ lookup table (LUT) on Fiji. The same contrast settings were used for each image pair. Scale bar = 5μm. (B) Bar graph showing quantitation of whole-cell GFP intensity from the Gnp1-GFP strains imaged in (A). Shown are standard error of the mean (SEM) and *p*-value from the student’s t-test demonstrating that the change observed is significant. (C) Depiction of the BirA-AviTag system where BirA-tagged Emc6 can biotinylate AviTag-EMC components (such as Emc2) and AviTag-substrates which it inserts into the membrane and only interacts with transiently. (D) Streptavidin blots of diploid strains expressing either AviTag-Fks1, -Gnp1, -Pdr5, -Pdr12 or -Stv1 (as negative control) together with BirA-Emc6, grown in the conditions described in Figure 1E. Blots showing AviTag-Fks1 and AviTag-Gnp1 are set to the same contrast, and the three remaining blots are set to the same higher contrast. The expected molecular weight in kDa for each tested protein including AviTag is written in parentheses after the protein name.

More broadly however, verifying transient interactions is, in itself, a challenging task as methods to validate PPIs (such as co-IP) are again optimized for very stable interactions. We therefore used a parallel biotinylation approach involving the BirA biotin ligase which specifically biotinylates the AviTag sequence (Cronan, 1990; Beckett *et al*, 1999). In this set- up, protein-client interactions can be assayed *in vivo* and at physiological expression levels. To do this, a haploid strain expressing BirA-Emc6 under its native promoter was mated with a haploid strain of the opposite mating-type which expressed the interactors N-terminally tagged with AviTag (also under native promoter control). The diploid strains were then analysed for the appearance of a streptavidin positive band that proves that BirA came sufficiently close to the AviTag (illustrated in Figure 3C). Initially, well-characterised and previously validated interactors were selected to test the utility and feasibility of this validation method: hence strains expressing either AviTag-Emc2, -Emc4, or -Spf1 (Jonikas *et al*, 2009; Shurtleff *et al*, 2018) were crossed with the BirA-Emc6 strain. In addition, the ABOLISH system was integrated into these strains to elucidate whether endogenous biotinylation reduction also confers any advantage in this system (Figure EV3A). Although the bands corresponding to biotinylated AviTag-Emc2, -Emc4, and -Spf1 were easy to distinguish from those of endogenously biotinylated proteins, auxin treatment very clearly reduced this background signal and made the assay even cleaner (Figure EV3B). This becomes critically important for lower abundance proteins, more transient interactions, proteins that are less efficiently biotinylated, or proteins whose molecular weight is similar to that of endogenously biotinylated proteins. Hence ABOLISH can also extend the dynamic range of BirA-AviTag for pairwise, gel-based assays.

Next, we investigated whether this method could be used to detect transient interactions between the EMC and its clients. Indeed, Fks1, an interactor found by TurboID (Figure 2B) and a known EMC substrate (Shurtleff *et al*, 2018), was readily detected by streptavidin blot using the BirA-AviTag/ABOLISH system (Figure 3D, left-most panel). The AviTagged amino acid permease, Gnp1, was similarly easy to detect. Some clients required a higher contrast setting to be visualised, however both AviTag-Pdr5 and AviTag-Pdr12 produced clear streptavidin-reactive bands compared to the negative control, AviTag-Stv1, which did not produce a detectable Emc6 interaction (Figure 3D). Collectively, these data highlight the power of the BirA-AviTag/ABOLISH system for providing a rare, *in vivo* ‘snapshot’ of the transient interactions between both previously-confirmed (Shurtleff *et al*, 2018; Bai *et al*, 2020) and newly-validated (Figure 3A, B and D) substrates and their insertase, the EMC. More broadly it serves as a rapid, systematic evaluation of TurboID interactomes.

### Generating a biotinylation toolkit: a collection of five full-genome libraries to facilitate high-throughput protein-protein interaction discovery

Through studying and comparing the promiscuous biotin ligases that can be used in yeast, we have demonstrated that TurboID, especially when combined with the ABOLISH system, serves as an unbiased tool to efficiently label several stable and transient functional interactors in *S.cerevisiae*. This in turn can lead to the discovery of novel protein-machinery substrates, as highlighted for the EMC (Figures 2 and 3) as well as regulators. We have also demonstrated the suitability of the BirA-AviTag technology for assaying and validating native pairwise interactions and the capacity of this sensitive methodology to highlight even transient interactions.

To truly harness the power of these biotinylation tools and make them widely applicable, we created whole-proteome collections of yeast strains (also called libraries) using our recently developed approach for yeast library generation called SWAp Tag (SWAT) (Yofe *et al*, 2016; Weill *et al*, 2018; Meurer *et al*, 2018). This approach allows us to take an initial library and swap its tag to any one of our choice in an easy and rapid manner. Therefore, using the N’ GFP SWAT library and accompanying SWAT protocol (Yofe *et al*, 2016; Weill *et al*, 2018) we generated five whole-genome libraries (Figure 4). In the first two, each strain encodes one yeast protein fused at its N’ to a TurboID-HA tag expressed under the control of a medium-strength constitutive *CYC1* promotor and generic N’ localisation signals (signal peptides (SPs) and mitochondrial targeting signals (MTS), (Yofe *et al*, 2016; Weill *et al*, 2018)) with, or without, the ABOLISH system. The third library is an N’ tag *CYC1*pr-BioID2-HA collection.

**Figure 4:**
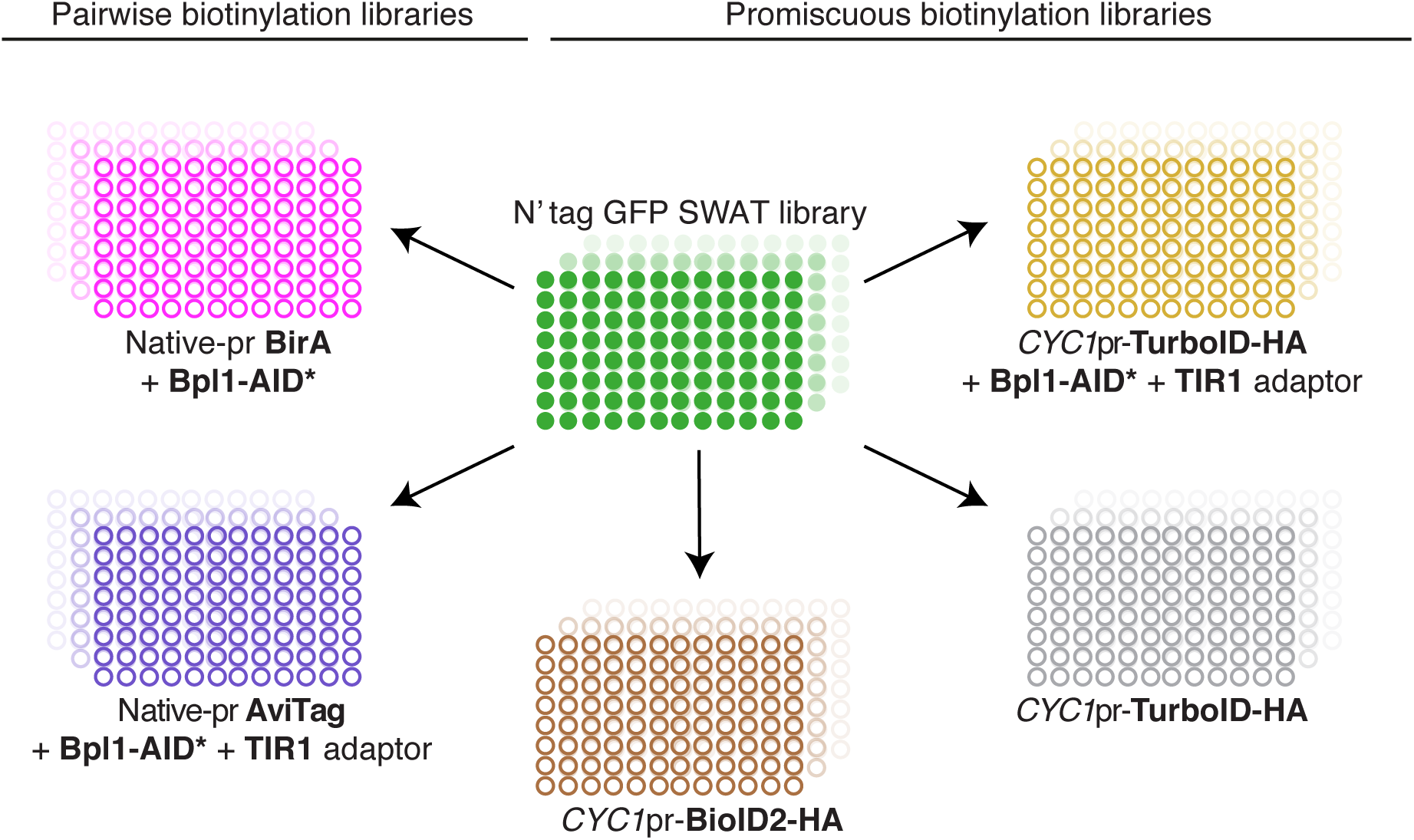
Generating five full-genome libraries using the SWAT method. Schematic representation of the creation of new library collections. The original N’ tag GFP SWAT library was used to generate five full-genome libraries using an automated process of mating, selection, sporulation and SWATing (see Methods for details). All promiscuous biotin ligase libraries have a *CYC1*pr upstream of the BioID2-HA/TurboID-HA, whereas the pairwise biotinylation libraries have BirA and AviTag downstream of the native promoter and targeting sequences. The AviTag/ABOLISH and TurboID-HA/ABOLISH libraries express the OsTIR1 adaptor protein and Bpl1-AID*-6HA or Bpl1-AID*-9myc, respectively. The BirA library includes only Bpl1-AID*-9myc.

The last two full-genome libraries express N-terminally tagged proteins with either BirA or AviTag under their endogenous promoters and N’ localisation signals, and the ABOLISH system is also integrated. In addition to PPI validation (as shown above) these libraries can, of course, be used for hypothesis-driven interrogation of interactions between any two proteins of interest.

All newly-generated libraries were subject to strict quality control checks (see Methods). Furthermore, a number of new library strains were selected and subject to SDS-PAGE analysis to confirm both protein expression and that the new tag had recombined in-frame during the SWAT process. This was demonstrated to be the case for the BioID2-HA (Figure EV4A), TurboID-HA (Figure EV4B) and TurboID-HA/ABOLISH (Figure EV4C) libraries. Hence these represent five high-coverage yeast libraries that will be freely distributed to enable high-throughput exploration, discovery and validation of stable and transient interactions throughout the yeast proteome.

## Discussion

Our work demonstrates the power of proximity biotin labelling tools for the exploration of both stable and transient protein interaction in yeast. We also show, for the first time, the utility of TurboID in this model organism. Surprisingly, previously-utilised protocols of growth in low biotin media to reduce endogenous biotinylation levels (Jan *et al*, 2014) also resulted in the downregulation of TurboID-tagged proteins. This demonstrates the advantage of using our ABOLISH system, which is designed to increase streptavidin-specific signal-to-noise through controlled Bpl1 degradation. Indeed, more interactors were found by TurboID when it was coupled to the ABOLISH system. Furthermore, the ABOLISH approach should be straightforward to extend to human cells since they also harbour only a single native biotin ligase (Uniprot ID: P50747).

The multi-subunit EMC (Jonikas *et al*, 2009; Christianson *et al*, 2012) has recently been characterised as an ER membrane insertase (Guna *et al*, 2018), and as such several substrates have been elucidated, particularly in human cells (Guna *et al*, 2018; Shurtleff *et al*, 2018; Chitwood *et al*, 2018; Tian *et al*, 2019; Volkmar *et al*, 2019; O’Keefe *et al*, 2021; Leznicki *et al*, 2021). Using EMC as a test case for TurboID utility in baker’s yeast, we discovered six new putative substrates of this complex, in addition to ones previously identified. In the past, EMC substrates were uncovered using *in vitro* assays (Guna *et al*, 2018; Chitwood *et al*, 2018; O’Keefe *et al*, 2021; Leznicki *et al*, 2021), labour-intensive ribosome profiling (Shurtleff *et al*, 2018), or proteomic profiling comparing control vs ΔEMC cells (Shurtleff *et al*, 2018; Tian *et al*, 2019; Volkmar *et al*, 2019; Bai *et al*, 2020). While loss-of-function studies have clearly proven useful, they suffer from both false negatives (from the presence of back-up systems (Ihmels *et al*, 2007)) and false positives (resulting from off-target effects). Endogenous labelling of transiently interacting substrates *in vivo* can therefore offer a more native approach to protein substrate discovery.

Of the six new candidate yeast EMC substrates that we identified, Alg1 stands out as unique. It is a highly-conserved and essential mannosyltransferase localised to LDs (Krahmer *et al*, 2013) and possesses a single N-terminal hydrophobic TMD. This type of substrate was only recently established to require the EMC for its biogenesis in humans (Leznicki *et al*, 2021). Notably, the free energy difference (ΔG, (Hessa *et al*, 2007)) for the TMD of Alg1 is -2.097, highly consistent with the ΔG values observed for the TMDs of human EMC-dependent LD proteins (Leznicki *et al*, 2021).

In addition to Alg1, we also found that Gnp1 and Pdr12 behave as substrates. Both Gnp1 and Pdr12 were previously flagged as putative yeast EMC substrates (Shurtleff *et al*, 2018; Bai *et al*, 2020) however remained unvalidated. Additionally, it seems that EMC-dependence for these proteins is conserved throughout evolution, with the levels of SLC7A1 and ABCA3 (human homologs of Gnp1 and Pdr12, respectively, (Fenech *et al*, 2020)) reduced in EMC KO cells (Tian *et al*, 2019; Tang *et al*, 2017). Interestingly, a physical association between EMC3 and ABCA3 was also reported (Tang *et al*, 2017), supporting our evidence for an EMC-Pdr12 interaction.

Naturally, not all putative substrates were identified or confirmed using proximity biotinylation methods. There are several aspects of each method that should therefore be thought of when choosing which of the libraries to utilise for PPI detection. For example, BirA specifically biotinylates AviTag and both modules must be proximal in space and on the same side of the membrane for biotinylation to occur. TurboID on the other hand, has more labelling opportunities as it can biotinylate any topologically available lysine residue within an interactor sequence. Other differences to bear in mind include the fact that the BirA-AviTag assay is carried out in diploid cells; unlike the haploid TurboID-expressing strains. Also, the TurboID-tagged proteins are under control of the constitutive *CYC1* promoter and generic N’ localisation signals. This is in contrast to the proteins tagged with BirA/AviTag which are under control of their native promoter and localisation signals. These native features provide much more physiological conditions, even though some low abundance proteins may be less easy to detect. Finally, as opposed to proteomic-based approaches, the BirA-AviTag system serves as a much cheaper and easy-to-use method requiring no specialist equipment.

In addition, despite the clear benefits of the ABOLISH system, we generated a TurboID ‘only’ library for instances where the addition of auxin may interfere with the proteins being studied, such as for TORC1 and its associated signalling pathways (Nicastro *et al*, 2021). Similarly, a BioID2 library was also included as part of our toolkit since it has already been adopted by the yeast community (Opitz *et al*, 2017; Singh *et al*, 2020). It is a smaller ‘tag’ compared to TurboID (27kDa vs 35kDa) and is known to have higher activity at higher temperatures; thus may prove useful for heat-shock experiments, for example.

We believe that the unique properties of the promiscuous and pairwise biotinylation machineries make the combination of both approaches the most powerful tool for stable, and even more so, transient PPI discovery and validation in baker’s yeast. Our whole-genome libraries and accompanying sample preparation protocols provide a broad resource for functional proteome exploration. All together, we present a complete biotinylation toolkit to enable high-throughput interaction profiling in baker’s yeast and the characterisation of new protein functions, signalling pathways and dynamic organellar processes that span the entire yeast proteome.

## Materials and methods

### Yeast strains and plasmids

All yeast strains used in this study are listed in Table S2. Strains were constructed using the lithium acetate-based transformation protocol (Gietz & Woods, 2002). All plasmids used are listed in Table S3 (see also (Longtine *et al*, 1998; Janke *et al*, 2004)) and primers were designed with the Primers-4-Yeast web tool (https://www.eizann.a.il/Primers-4-Yeast/, (Yofe & Schuldiner, 2014), Table S4). The original SWAT donor strain (yMS2085; (Weill *et al*, 2018) was transformed with SWAT donor plasmids encoding BioID2-HA or TurboID-HA. SWAT donor strains encoding the ABOLISH system were constructed by C-terminally tagging Bpl1 with AID*-9myc and integrating the OsTIR1 adaptor into the *HIS* locus using a PmeI-cut, OsTIR1-encoding plasmid (Morawska & Ulrich, 2013; Orgil *et al*, 2015). These strains were transformed with SWAT donor plasmids encoding TurboID-HA, BirA or AviTag.

### Yeast growth

Yeast cells were grown on solid media containing 2.2% agar or liquid media. YPD (2% peptone, 1% yeast extract, 2% glucose) was used for cell growth if only antibiotic selections were required, whereas synthetic minimal media (SD; 0.67% [w/v] yeast nitrogen base (YNB) without amino acids and with ammonium sulphate or 0.17% [w/v] YNB without amino acids and with monosodium glutamate, 2% [w/v] glucose, supplemented with required amino acid) was used for auxotrophic selection. Antibiotic concentrations were as follows: nourseothricin (NAT, Jena Biosciences) at 0.2g/L; G418 (Formedium) at 0.5g/L; and hygromycin (HYG, Formedium) at 0.5g/L. Yeast grown for transformation, protein extraction or LC-MS/MS analysis were first grown in liquid media with full selections overnight at 30°C and subsequently back-diluted into YPD/SD media to an OD_600_ of ∼0.2. Cells were collected after at least one division but before reaching an OD_600_ of 1 and either immediately used for transformation or snap frozen for later processing. Cells grown for streptavidin-AP followed by LC-MS/MS were treated for ∼18h with 50μM biotin as in (Roux *et al*, 2012; Branon *et al*, 2018; Larochelle *et al*, 2019; Singh *et al*, 2020). For all ABOLISH experiments, biotin was used at 100nM with the exception of Figure EV1B, where it was used at 10nM (Jan *et al*, 2014). For figures EV1E, EV1D and EV1E, RB media was prepared as specified in (Jan *et al*, 2014). Auxin was used at 1mM. Times of treatments are specified in the appropriate figure legends.

### Protein extraction and SDS-PAGE analysis

Cells pellets were resuspended in 200μl lysis buffer (8M urea, 50mM Tris pH 7.5, protease inhibitors (Merck)). From there on, processing, SDS-PAGE separation, Western blotting, and fluorescent-based imaging was done as described in (Eisenberg-Bord *et al*, 2021), with the exception of the HA blot in Figure EV1C. Here, the SDS-PAGE gel was blotted onto PVDF membrane (Millipore) by wet transfer and imaging was done using X-ray film (FujiFilm) to detect signal from HRP-conjugated anti-mouse secondary antibody (1:7500, Jackson ImmunoResearch, #111-035-003) incubated with ECL substrate (Thermo Scientific). The following antibodies were used for Western blot: anti-HA (1:1000, BioLegend, #901502), anti-myc (1:3000, Abcam, #ab9106), anti-Histone H3 (1:5000, Abcam, #ab1791), anti-Sec61 (1:5000, a kind gift from Matthias Seedorf of Heidelberg University and Marius Lemberg of the University of Cologne), anti-Actin (1:2000, Abcam, #ab8224), goat anti-rabbit IgG H&L 800CW (1:7500, Abcam, #ab216773), goat anti-mouse IgG H&L 680RD (1:7500, Abcam, #ab216776). Membranes were incubated for 1h at RT with fluorescent streptavidin (1:10000, Invitrogen, #S11378) diluted in 2% (w/v) BSA/PBS containing 0.01% NaN_3_ to detect biotinylated proteins.

### Immunoprecipitation and LC-MS/MS sample preparation

Cell pellets from a total of ∼5ODs were resuspended in 400μl lysis buffer (150 mM NaCl, 50mM Tris-HCl pH 8.0, 5% Glycerol, 1% digitonin (Sigma, #D141), 1mM MgCl_2_, protease inhibitors (Merck), benzonase (Sigma, #E1014)). The cell suspension was then transferred to a 2ml FastPrep™ tube (lysing matrix C, MP Biomedicals) and lysis was carried out by 6 x 1min maximum speed cycles on a FastPrep-24™ cell homogeniser (MP Biomedicals), with the samples being returned to ice for 5min between each cycle. Lysates were cleared at 16,000g for 10min at 16,000g at 4°C and the supernatant was transferred to a fresh microcentrifuge tube. For HA-IP, samples were first incubated for 1h at 4°C with 2μl of anti-HA antibody (BioLegend) and then for another hr after adding 30μl of washed magnetic ProteinG beads (Cytiva). The beads were washed twice with 200μl of digitonin wash buffer (150mM NaCl, 50mM Tris-HCl pH 8.0, 1% digitonin) and then four times in basic wash buffer (150mM NaCl, 50mM Tris-HCl pH 8.0) before being incubated with 50μl elution buffer (2M urea, 20mM Tris-HCl pH 8.0, 2mM DTT, and 0.5μl trypsin (0.5μg/μl, Promega, #V5111)) per sample for 90min. The eluate was removed from the beads and collected in a fresh microcentrifuge tube. 50μl alkylation buffer (2M urea, 20mM Tris-HCl pH8.0, 50mM iodacetamide (IAA)) was then added to the beads and incubated for 10min. This buffer was also removed from the beads and combined with the first eluate. Finally, the beads were washed with 50μl urea buffer (2 M urea, 20 mM Tris-HCl pH 8.0) for another 10min, and again the buffer was removed and combined with the above mixture. All elution steps were carried out at room temperature (RT) in the dark with shaking (1400rpm). The eluted mixture (150μl total volume) was incubated overnight at RT in the dark at 800rpm. The following morning 1μl 0.25μg/μl trypsin was added to each sample and incubated for a further 4h at RT in the dark at 800rpm. For streptavidin-AP, the same protocol as for HA-IP was used with the following changes: (1) 100μl streptavidin-conjugated beads (Cytiva) were used per sample and incubated overnight at 4°C; (2) post-AP beads were washed twice in 500μl 2% SDS wash buffer (2% v/v SDS (BioRad, #1610418), 150mM NaCl, 50mM Tris-HCl pH 8.0), twice in 500μl 0.1% SDS wash buffer, then twice in 500 μl basic wash buffer. All washes were 5min and were carried out at RT on overhead rotator. Following digestion, peptides were desalted using Oasis HLB, μElution format (Waters, Milford, MA, USA). The samples were vacuum dried and stored at -80°C until further analysis.

### LC-MS/MS settings and analysis

ULC/MS-grade solvents were used for all chromatographic steps. Each sample was loaded using split-less nano-Ultra Performance Liquid Chromatography (10kpsi nanoAcquity; Waters, Milford, MA, USA). The mobile phase was: (A) H_2_O + 0.1% formic acid and (B) acetonitrile + 0.1% formic acid. Desalting of the samples was performed online using a reversed-phase Symmetry C18 trapping column (180μm internal diameter, 20mm length, 5μm particle size; Waters). The peptides were then separated using a T3 HSS nano-column (75μm internal diameter, 250mm length, 1.8μm particle size; Waters) at 0.35μL/min.

Peptides were eluted from the column into the mass spectrometer using the following gradient: 4% to 30% B over 55min, 30% to 90% B over 5min, maintained at 90% for 5min and then back to the initial conditions. The nanoUPLC was coupled online through a nanoESI emitter (10μm tip; New Objective, Woburn, MA, USA) to a quadrupole orbitrap mass spectrometer (Q Exactive HF; Thermo Scientific) using a FlexIon nanospray apparatus (Proxeon). Data was acquired in data dependent acquisition (DDA) mode, using a Top10 method. MS1 resolution was set to 120,000 (at 200m/z), mass range of 375-1650m/z, AGC of 3e6 and maximum injection time was set to 60 msec. MS2 resolution was set to 15,000, quadrupole isolation 1.7m/z, AGC of 1e5, dynamic exclusion of 20sec and maximum injection time of 60msec.

### LC-MS/MS raw data processing

Raw data was processed with MaxQuant v1.6.6.0. The data was searched with the Andromeda search engine against the SwissProt *S. cerevisiae ATCC204508/S288c* proteome database (November 2018 version, 6049 entries) in addition to the MaxQuant contaminants database. All parameters were kept as default except: Minimum peptide ratio was set to 1, maximum of 3 miscleavages were allowed, and match between runs was enabled. Carbamidomethylation of C was set as a fixed modification. Oxidation of M, deamidation of N and Q, and protein N-term acetylation were set as variable modifications. The LFQ intensities were used for further calculations using Perseus v1.6.2.3. Decoy hits were filtered out, as well as proteins that were identified on the basis of a modified peptide only. The LFQ intensities were log2-transformed and only proteins that had at least 2 valid values in at least one experimental group were kept. The remaining missing values were imputed by a random low range distribution. Student’s t-tests were performed between the relevant groups to identify significant changes in protein levels. The mass spectrometry proteomics data have been deposited to the ProteomeXchange Consortium via the PRIDE (Perez-Riverol *et al*, 2021) partner repository with the dataset identifier PXD033348.

### Imaging

Cells were grown overnight in 100μl SD media (with appropriate amino acid (AA) selections) in a round-bottomed 96-well plate (ThermoFisher) at 30°C with shaking. 5μl of overnight culture was back-diluted into 95μl YPD and incubated for ∼4h at 30°C with shaking. 50μl of culture was subsequently transferred to a Concanavalin A (ConA, Sigma, 0.25mg/ml)-coated 384-well glass-bottomed microscopy plate (Matrical Bioscience) and incubated for 20min at RT. The media was removed and the cells were washed twice in 50μl SD-riboflavin+completeAA prior to being imaged in the same media at RT. Images for the top and middle panels of Figure 3A were using a VisiScope Confocal Cell Explorer system (Visitron Systems) coupled to an inverted IX83 microscope (Olympus), a CSU-W1-T1 50μm spinning disk scanning unit (Yokogawa) and an Edge sCMOS camera (PCO) controlled by VisiView software (V3.2.0, Visitron Systems). A 60x oil objective was used (NA = 1.42, Olympus) together with a GFP filter EX470/40nm, EM525/50nm (Chroma) and a 100mW 488nm laser (Visitron Systems). Images for the bottom panel of Figure 3A were using a SpinSR system (Olympus) coupled to a CSUW1-T2SSR spinning disk scanning unit (Yokogawa) and an ORCA-Flash 4.0 CMOS camera (Hamamatsu). A 60x air objective was used (NA = 0.9, Olympus) together with a GFP filter EX470/40nm, EM525/50nm (Chroma) and a 100mW 488nm laser system (Coherent OBIS LX). All images are single-focal plane.

Fiji was used for image inspection and brightness adjustment (Schindelin *et al*, 2012). For quantitation (Figure 3B), the images were first processed using the ScanR Analysis software (V3.2.0.0, Olympus). Processing used a neural network virtual channel module to segment the image (transmitted channel only) to cells using the intensity object segmentation module. Next, noise and objects that were poorly segmented were removed based ontheir area and circularity factor, and for each cell the mean signal intensity in the 488 channel was measured. The following Python script was used for data analysis: https://github.com/Maya-Schuldiner-lab/Analysis-of-GFP-quantified-and-segmented-cells.

### Diploid strain generation

The BirA-Emc6 strain was grown on YPD supplemented with NAT and AviTagged interactors were grown on SD_MSG_ without histidine supplemented with HYG at 30°C overnight. Both strains were velveted onto a YPD plate and grown overnight at RT. The mated strains were then velveted onto SD_MSG_ plates without histidine supplemented with NAT and HYG and grown overnight at 30°C. This step was repeated once more to select for diploid strains containing the combination of desired traits.

### BirA/AviTag interaction assay

Diploid strains were grown overnight at 30°C in SD_MSG_ liquid media without histidine supplemented with NAT, HYG and auxin (1mM, Sigma). Strains were then back-diluted to 0.2 OD_600_ and incubated for 4 hours at 30°C in SD_MSG_ liquid media without histidine supplemented with NAT, HYG, auxin and biotin (100nM, Sigma). Cells were collected upon reaching 0.5 OD_600_ by centrifugation at 3,000g for 3min, washed once in double distilled water (DDW) and then processed for Western blotting (see above).

### Yeast library generation

SWAT library generation was performed as described (Weill *et al*, 2018). Briefly, a RoToR array pinning robot (Singer Instruments) was used to mate the parental N’ tag GFP SWAT library with the required donor strain (Table S2) and carry out the mating, sporulation and selection protocol to generate a haploid library selected for all the desired features (Tong & Boone, 2007). Growth of the library on YPGalactose (2% peptone, 1% yeast extract, 2% galactose) was used to induce SceI-mediated tag swapping, and subsequent growth on SD containing 5-fluoroorotic acid (5-FOA, USBiological) at 1g/L, and required metabolic and antibiotic selections was used to selected for strains which had successfully undergone the SWAT process. Information on library genotypes, mating types, swap-tag efficiency, percentage survival and other quality control checks can be found in Table S5.

## Data availability

Protein interaction IP/AP-MS data: PRIDE, PXD033348 (Reviewer account username; password: y3CJEJli).

## Acknowledgements

We thank Dr. Yury Bykov, Dr. Ofir Klein, Rosario Valenti and Sivan Arad from the Schuldiner lab for critical feedback of this manuscript. We are grateful to Dr. Yael Elbaz-Alon, Dr. Yoav Peleg and Prof. Itay Onn for plasmids, Prof. Matthias Seedorf and Prof. Marius Lemberg for reagents, and to Corine Katina (The de Botton Institute for Protein Profiling) for help with LC-MS/MS sample preparation. The project was supported by the European Research Council Consolidator Grant OnTarget 864068, an Israel Science Foundation grant (ISF 760/17) and an SFB 1190 grant from the Deutsche Forschungsgemeinschaft (DFG). Maya Schuldiner is an incumbent of the Dr. Gilbert Omenn and Martha Darling Professorial Chair in Molecular Genetics.

## Author contributions

E. Fenech and N. Cohen designed, performed and analysed experiments. M. Kupervaser ran LC-MS/MS samples and processed LC-MS/MS data. Z. Gazi quantified microscopy data. E. Fenech and M. Schuldiner wrote the manuscript which all authors read and provided feedback on. M. Schuldiner supervised the work and secured funding.

---

**Expanded view figure 1:**
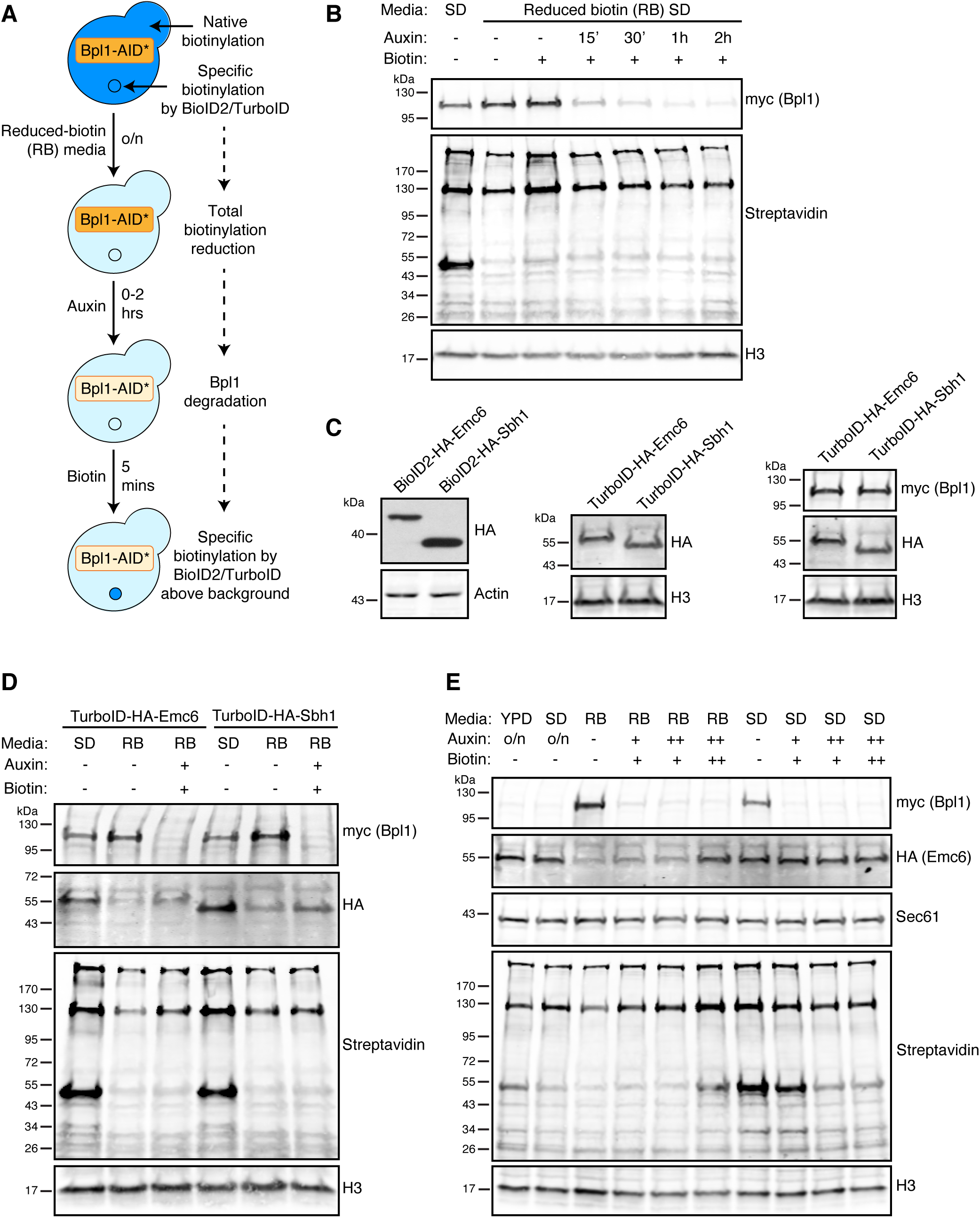
Optimising the conditions used for ABOLISH. (A) Schematic of the original growth conditions used to test Bpl1-AID*-9myc degradation and endogenous biotinylation reduction. (B) Anti-myc and streptavidin blots of cells expressing Bpl1-AID*- 9myc and OsTIR1 grown overnight and back-diluted in regular synthetic (SD) or reduced biotin (RB) media. To the cells grown in RB media, auxin was either omitted (-) or added 15min, 30min, 1hour or 2hours prior to harvesting. Similarly, biotin was either omitted (-) or added 5min prior to harvesting. (C) Anti-HA blots confirming the expression of BioID2/TurboID-HA-tagged Emc6 and Sbh1. An anti-myc blot was included for the strains containing the ABOLISH system. (D) Western blot analysis of cells expressing either TurboID-HA-Emc6 or TurboID-HA-Sbh1 together with the ABOLISH system. Cells were grown in SD or RB media as in (B). Auxin and biotin were either omitted (-) or added (+) 1.5hours and 30min (respectively) before harvesting. (E) Western blot analysis of cells expressing TurboID-HA-Emc6 and the ABOLISH system. Cells were grown in several different conditions: (i) overnight and back-diluted in rich media (YPD) containing auxin; (ii) overnight and back-diluted in regular SD containing auxin; (iii) overnight and back-diluted in RB media with auxin and biotin either omitted (-), added for 1.5hours and 30min (respectively) before harvesting (+), or added for the entire duration of the back-dilution (++); (iv) overnight and back-diluted in regular SD with auxin and biotin treatments as in (iii). Sec61 was used an untagged ER membrane protein control. For panels B-E, H3 (histone H3) or Actin were used as loading controls.

**Expanded view figure 3:**
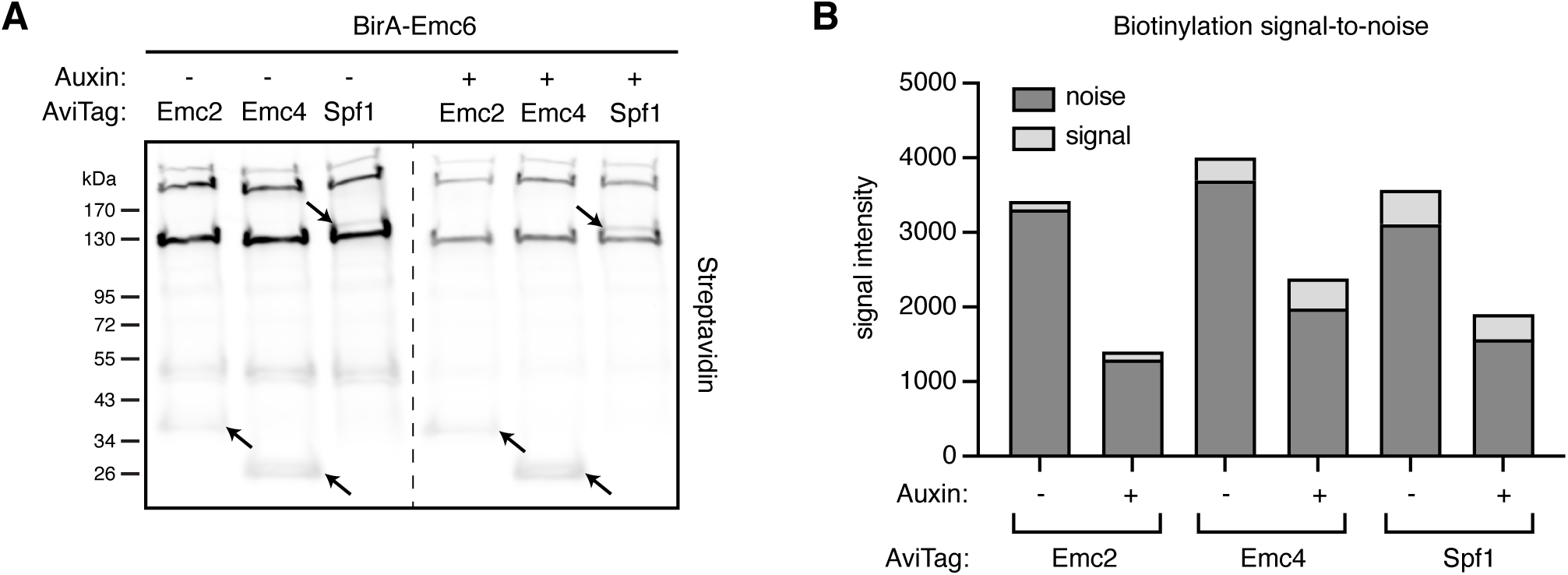
Contribution of the ABOLISH system to noise reduction in BirA- AviTag blotting. (A) Streptavidin blot of diploid strains expressing either AviTag-Emc2, - Emc4, or -Spf1 together with BirA-Emc6, grown in media with or without auxin. (B) Quantitation of the streptavidin signal from endogenously biotinylated proteins (noise) and biotinylated AviTagged-proteins (signal) shows a reduction of the background noise by ∼half when ABOLISH is activated.

**Expanded view figure 4:**
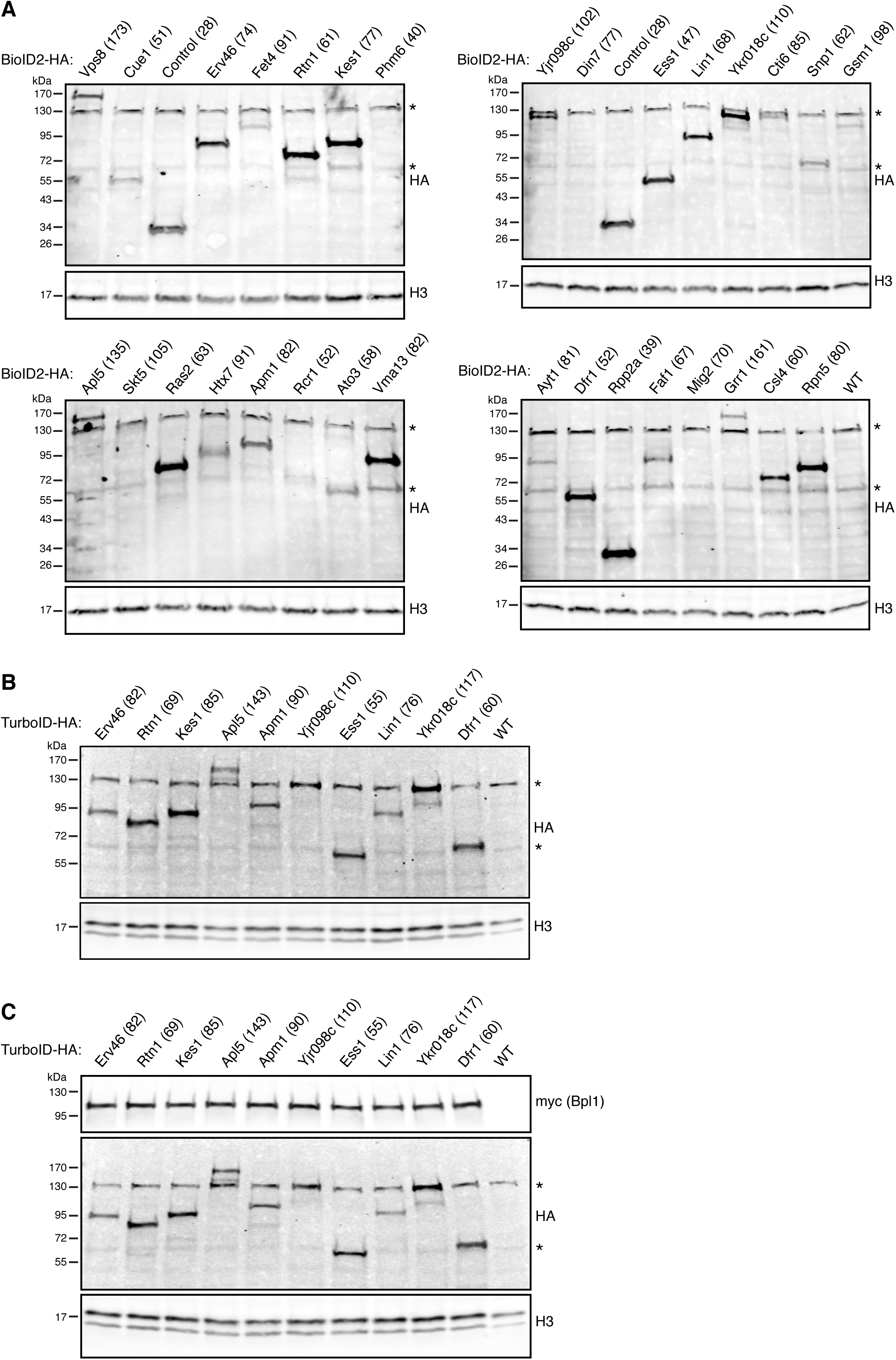
Quality control checks on biotinylation toolkit libraries. Western blot analysis for selected strains from the BioID2-HA (A), TurboID-HA (B) and TurboID- HA/ABOLISH (C) libraries. For the BioID2-HA library, Phm6, Din7, Skt5, Rcr1 and Mig2 all have low endogenous expression (relative intensity values of ≤ 27, (Weill *et al*, 2018)) which likely explains why they were not readily detectable. ‘Control (28)’ refers to BioID2-HA not tagged to any protein, and ‘WT’ denotes lysate ran from the BY4741 laboratory strain to highlight non-specific bands; the most prominent of which are marked with an asterisk. For all panels, H3 (histone H3) is used as a loading control. The expected molecular weight in kDa for each tested protein including their tag is written in parentheses after the protein name.

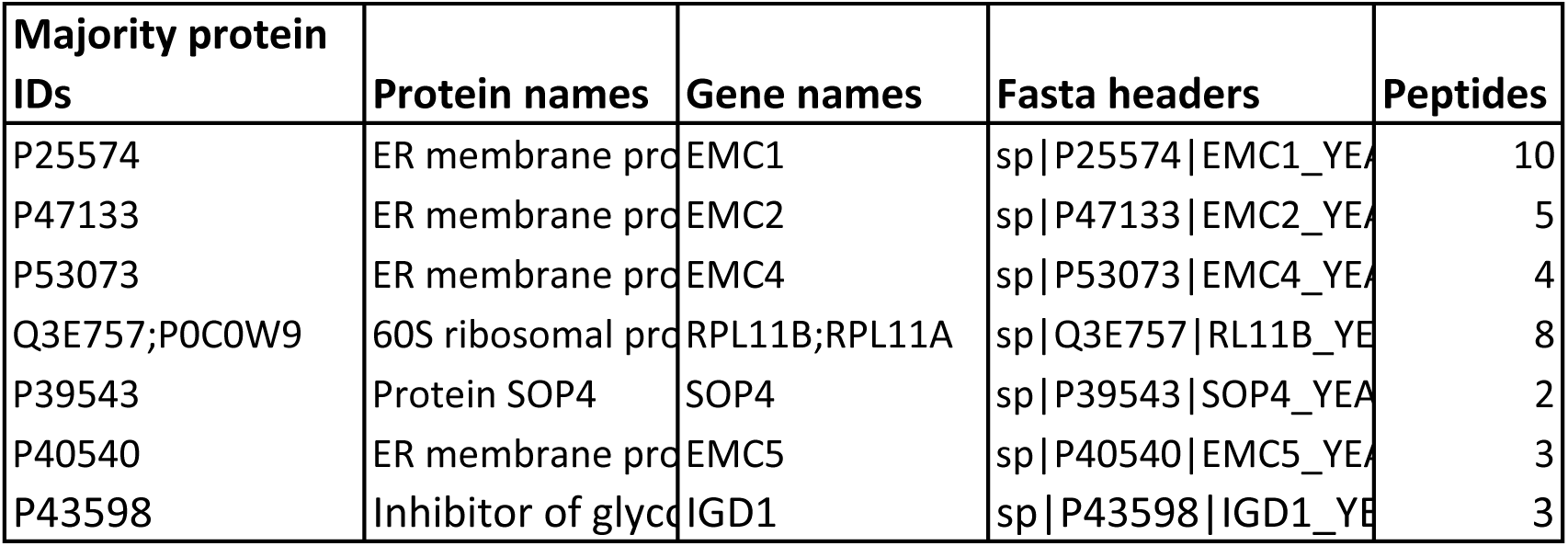
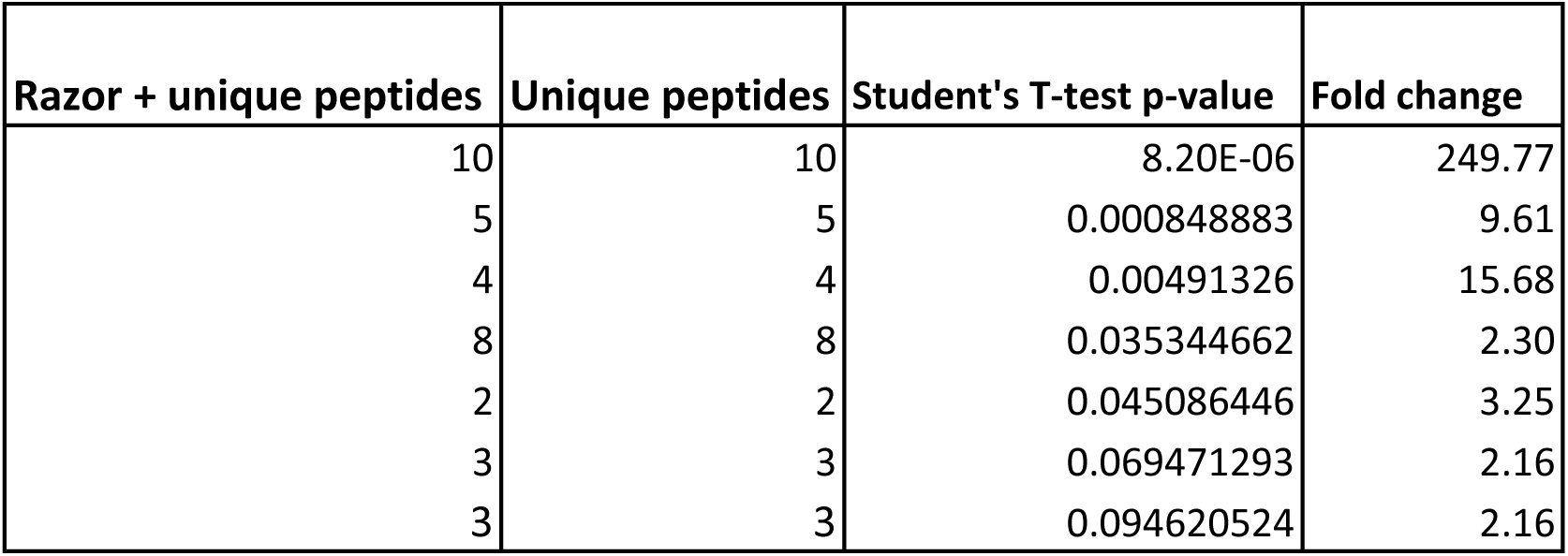
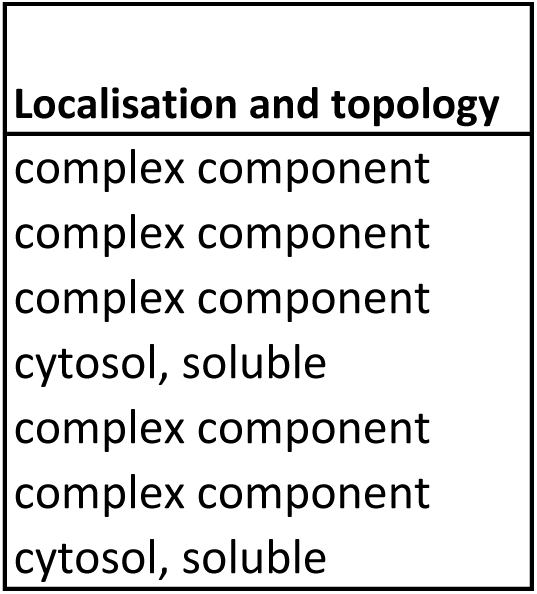

**Supplementary Table S2:**
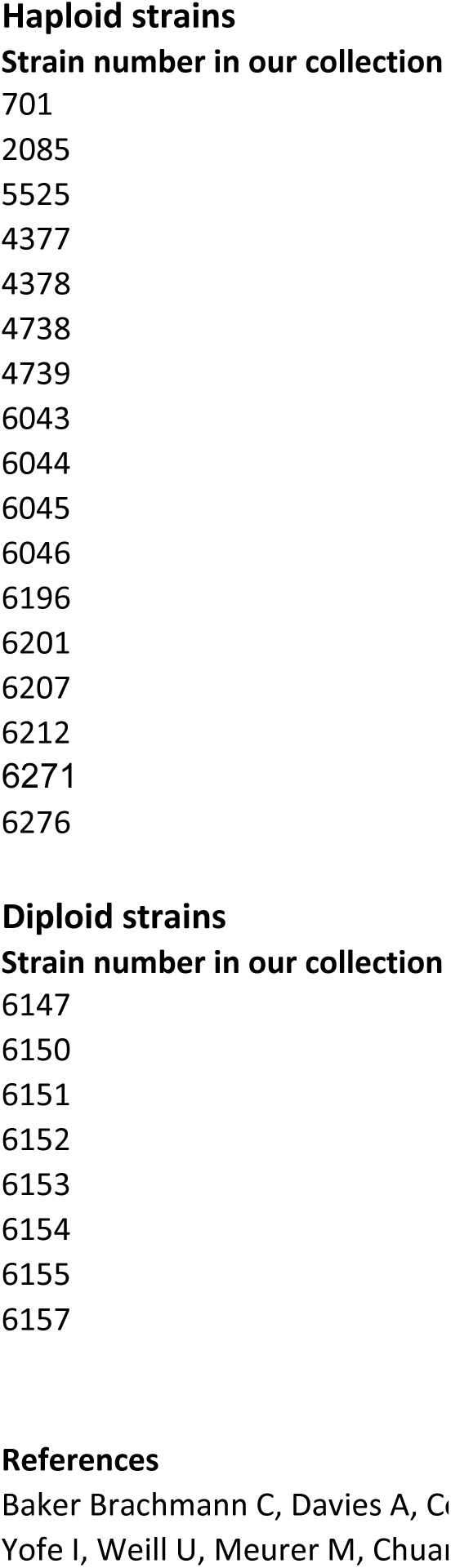
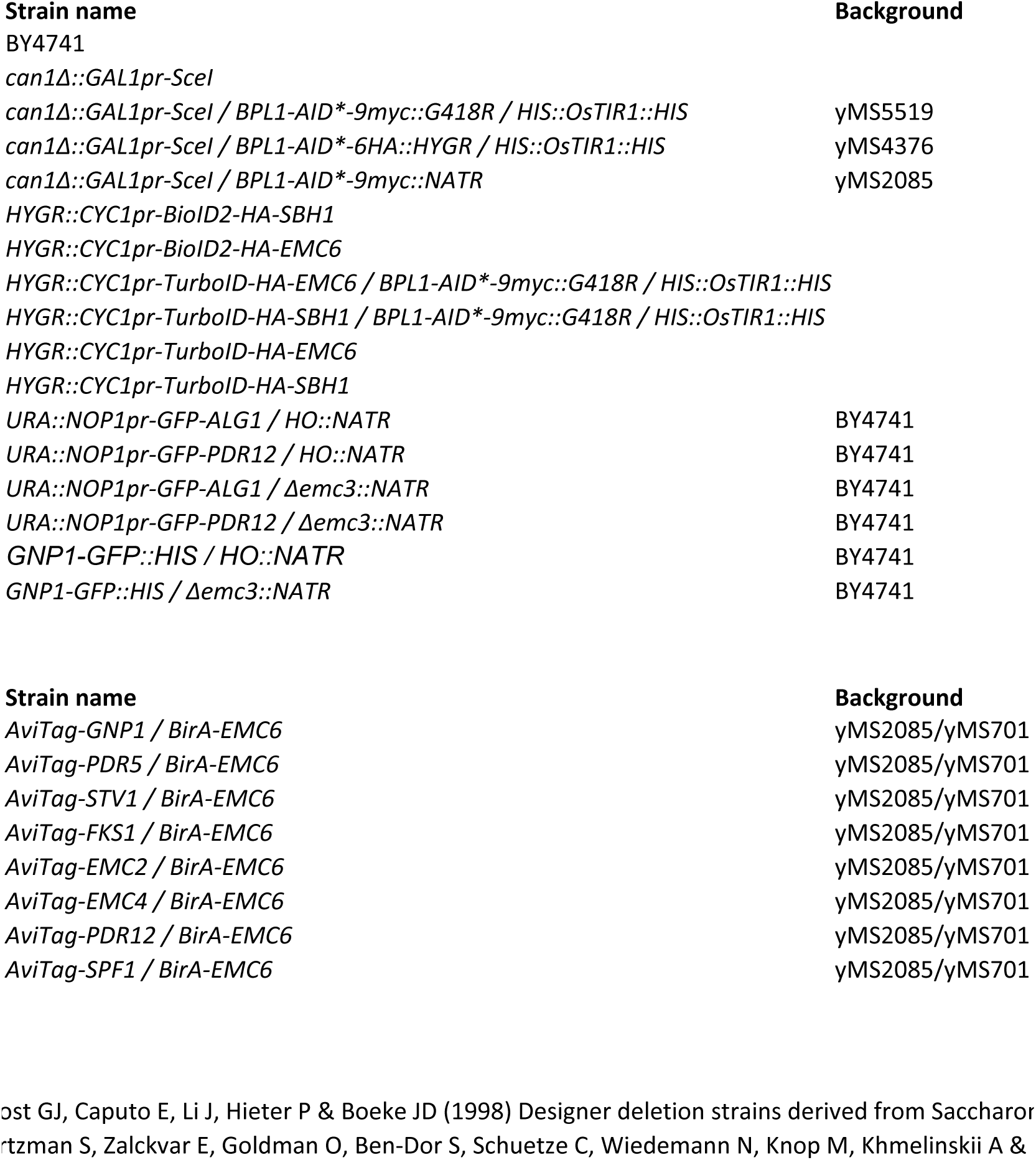
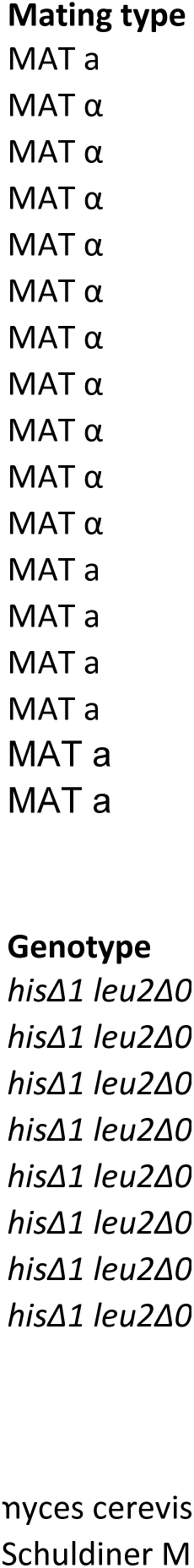
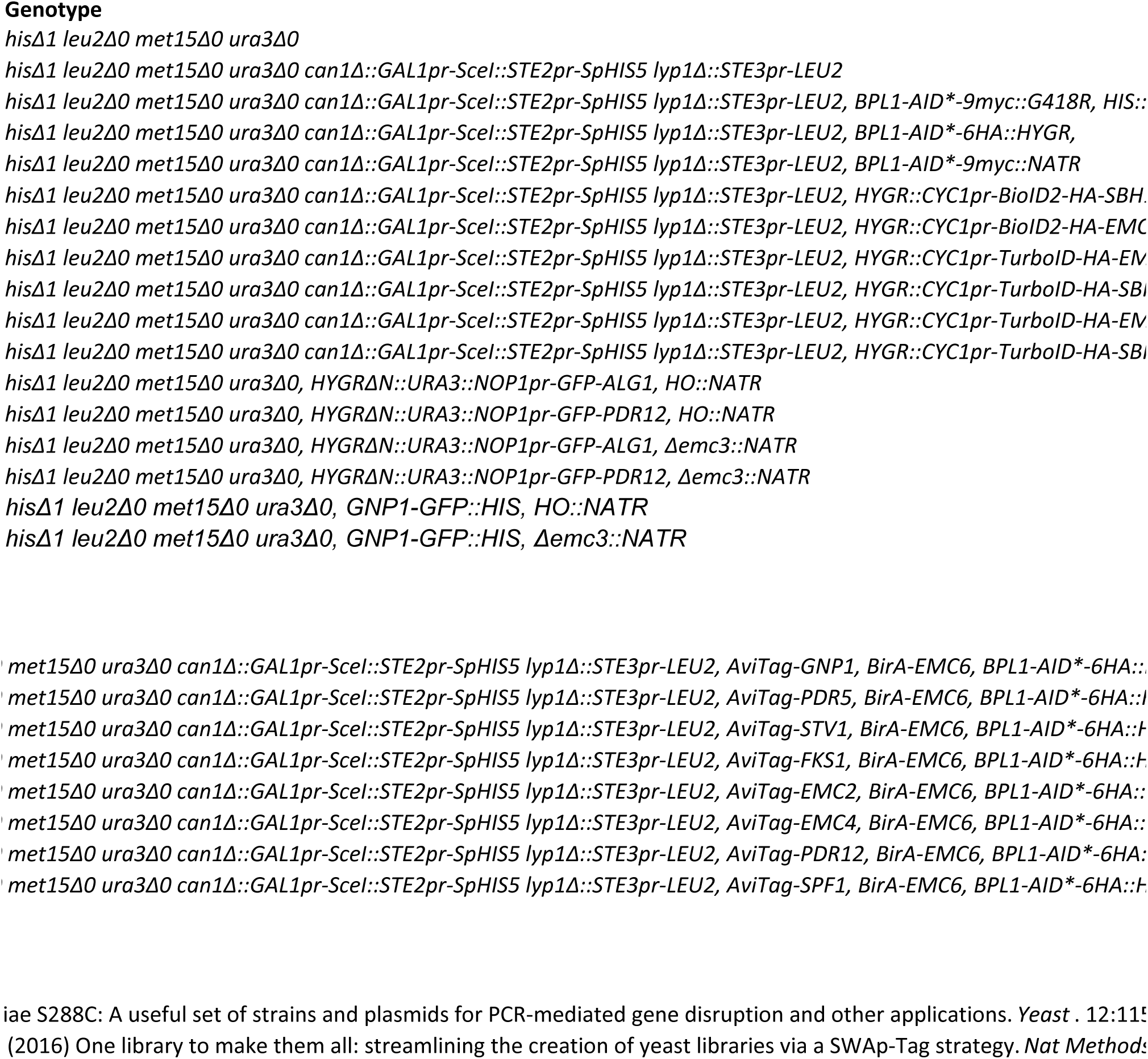
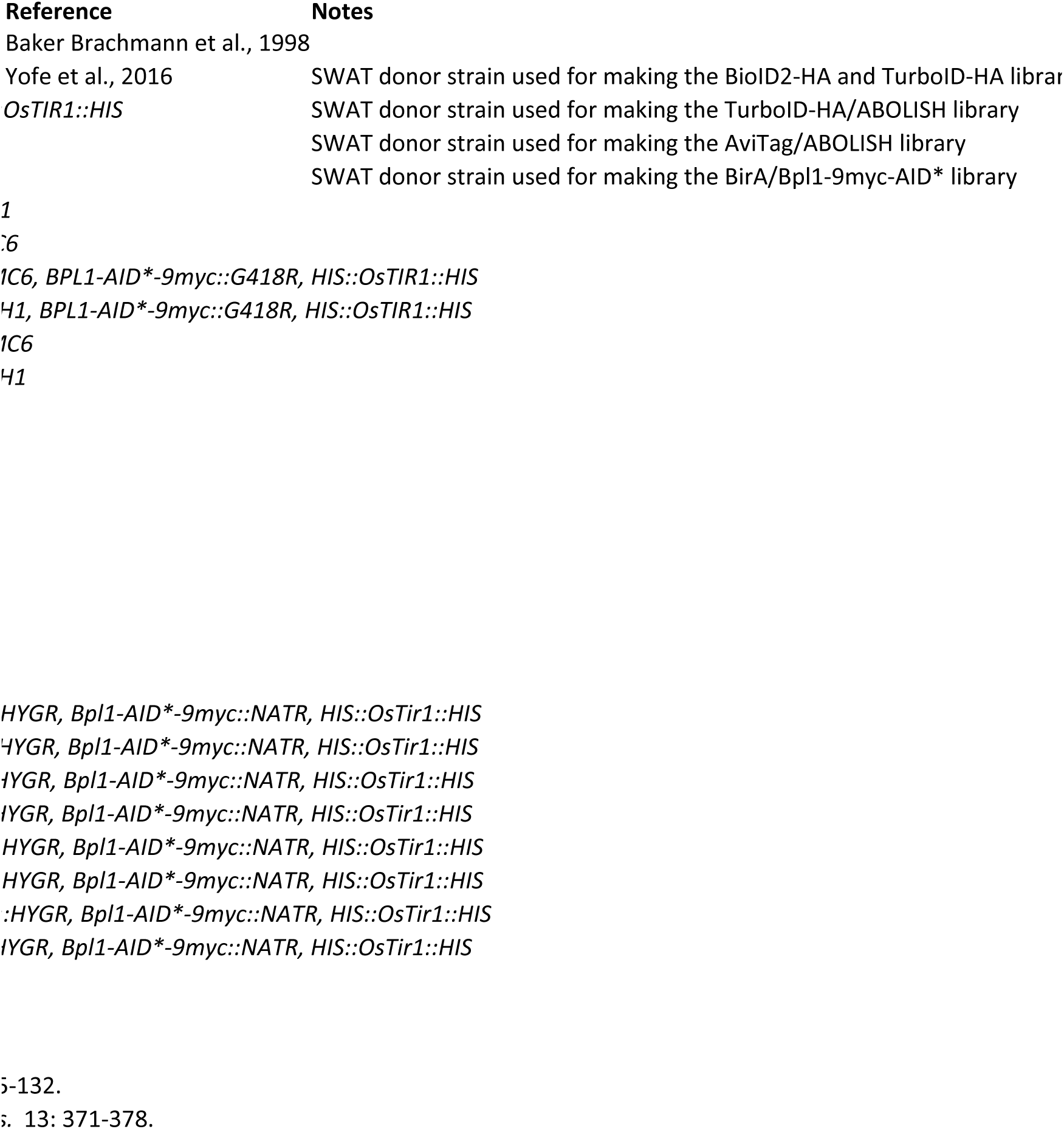
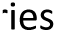
Yeast strains used in this study

**Supplementary Table 3:**
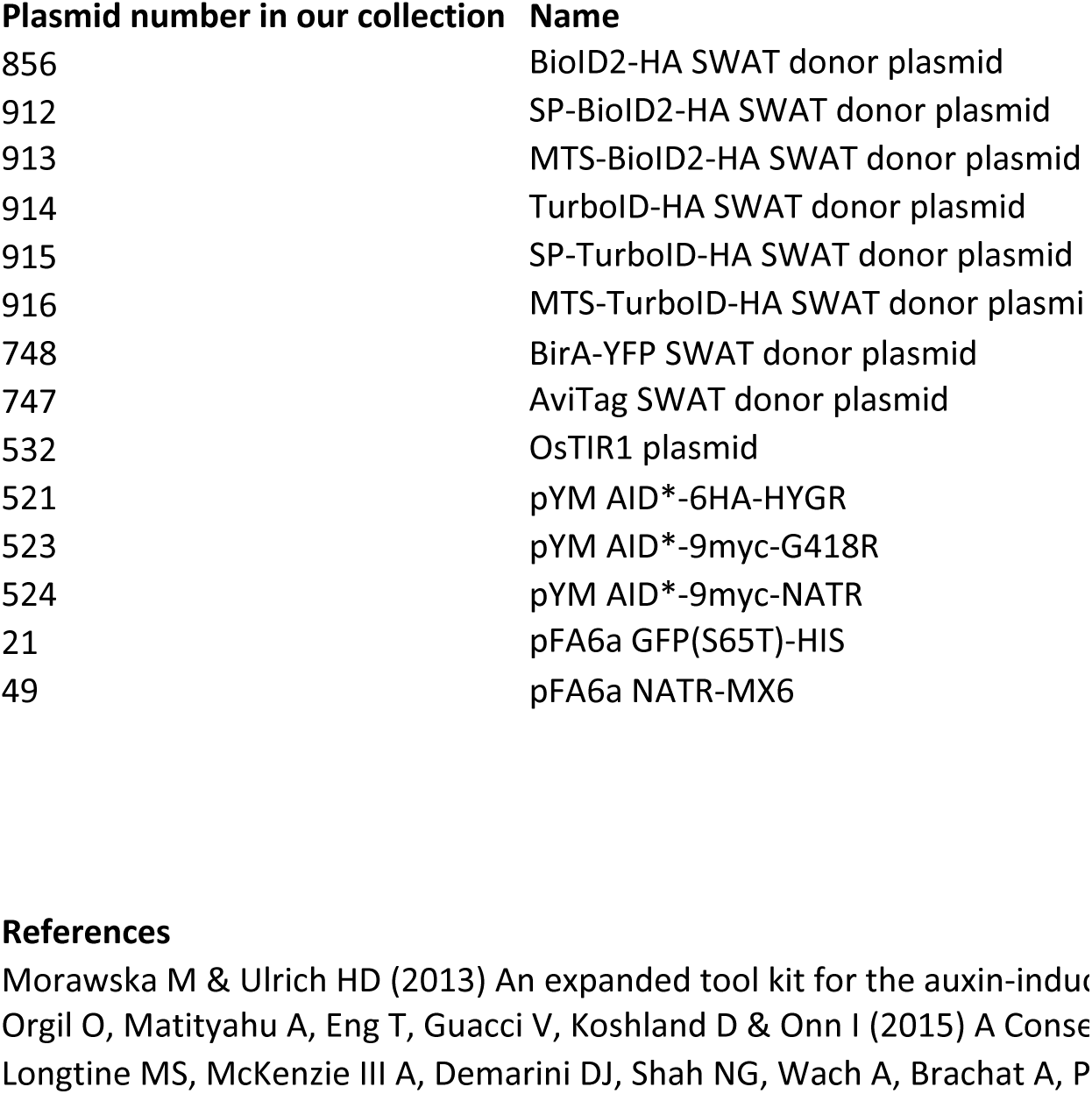
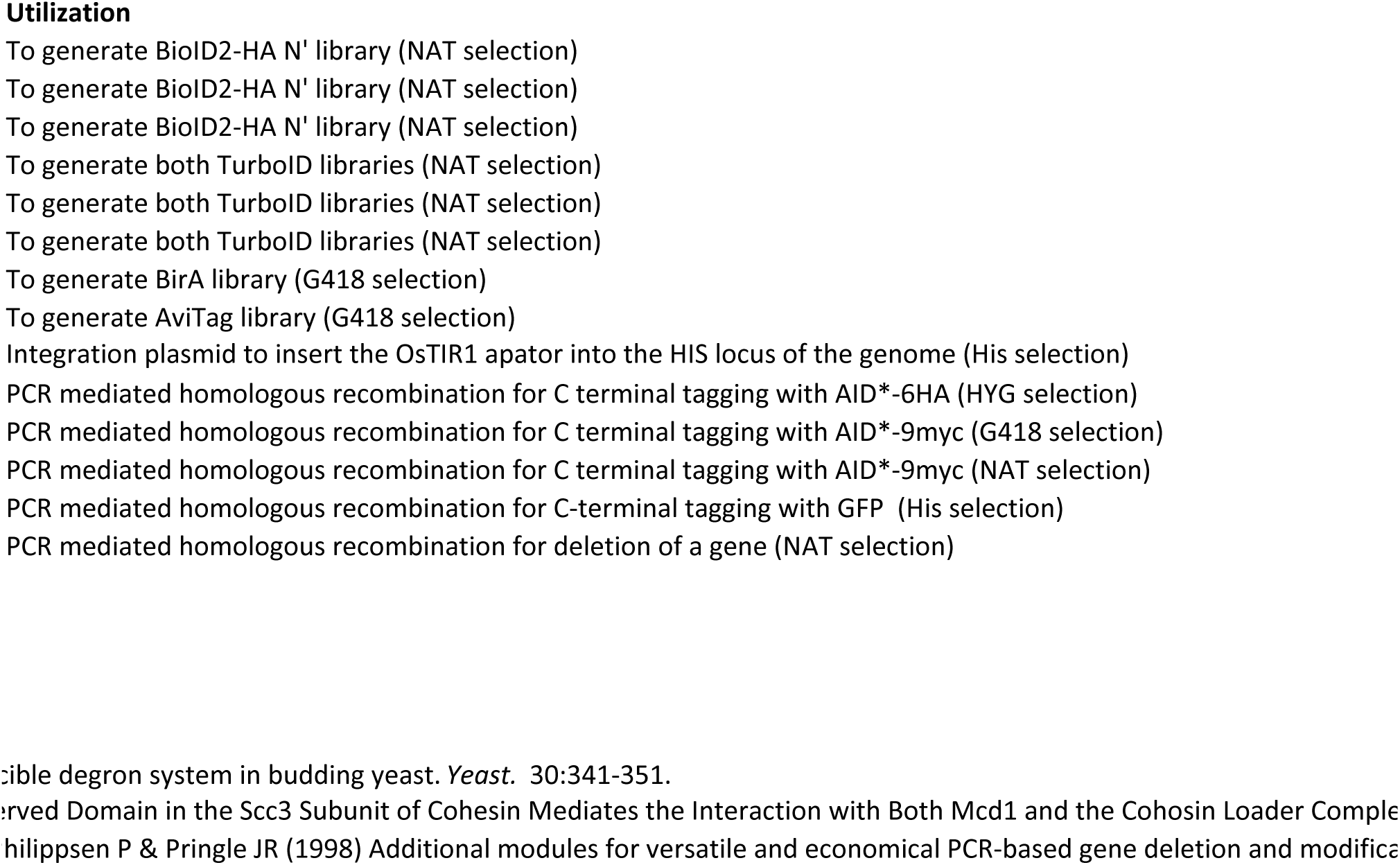
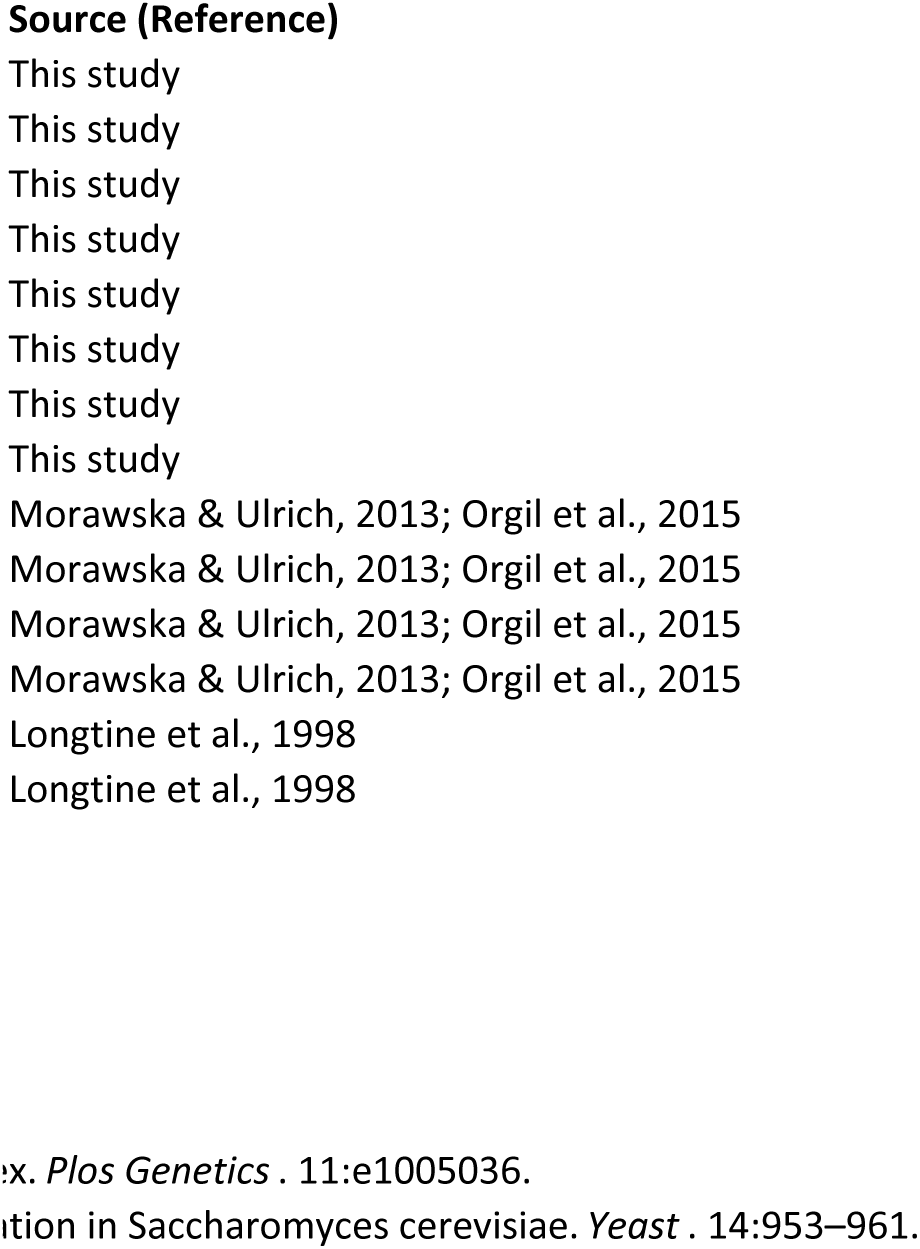
Plasmids used in this study

**Supplementary Table 4:**
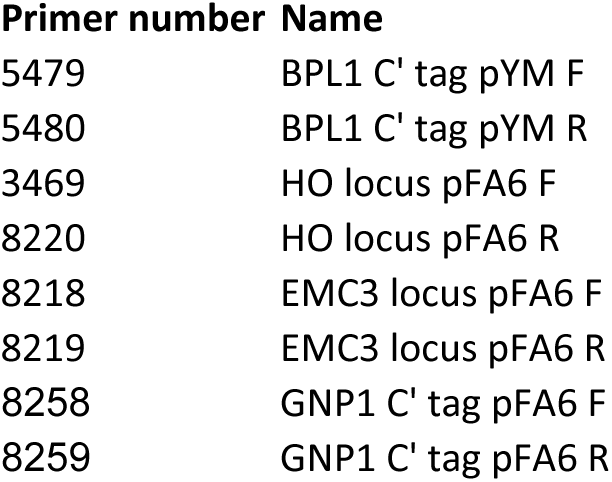
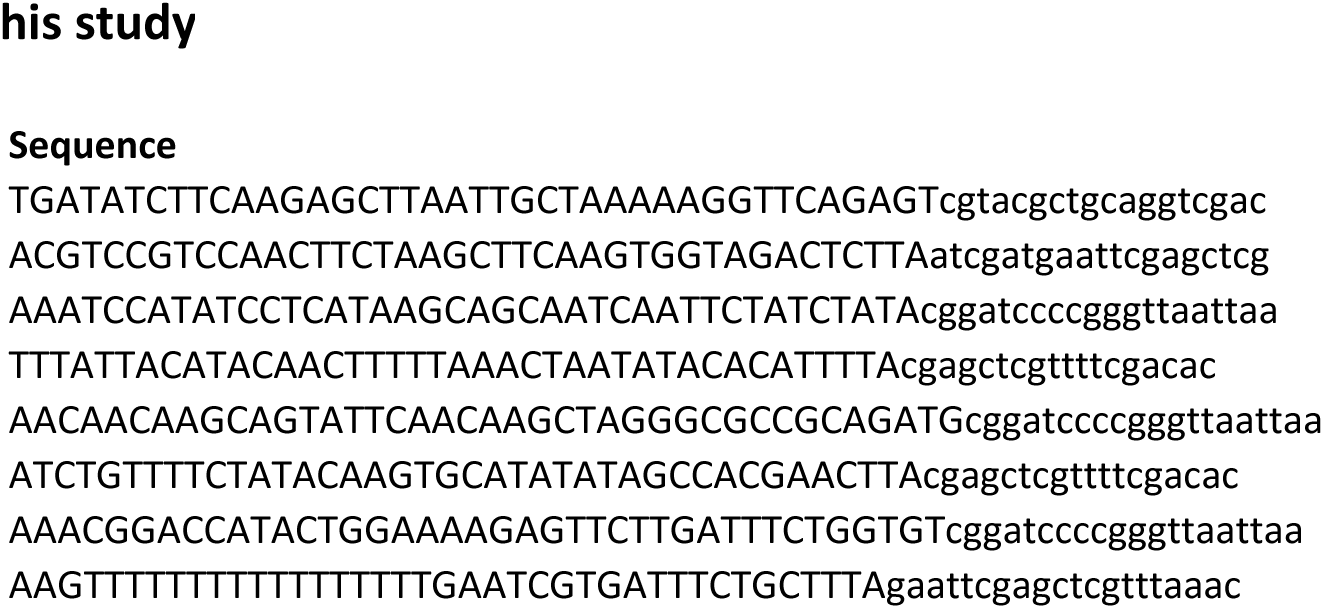
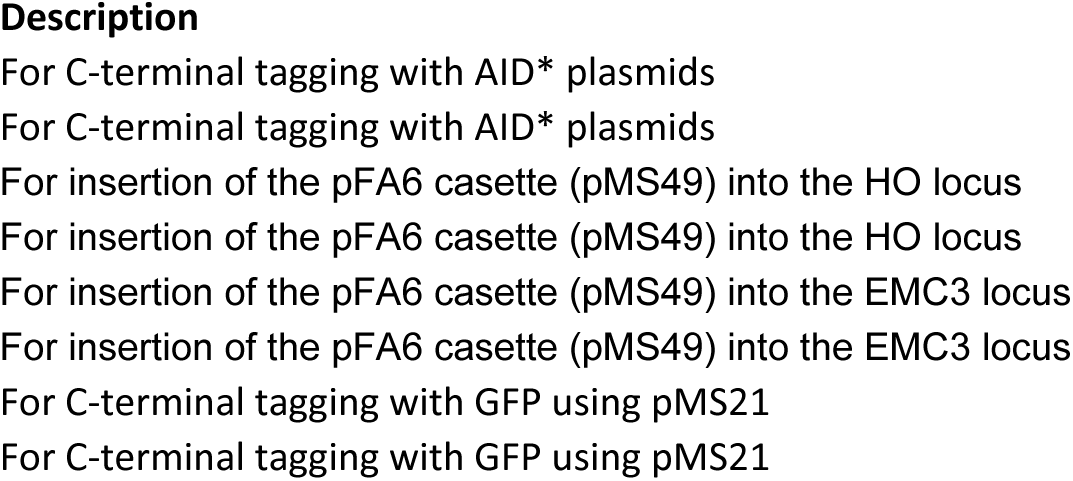
Primers used in t**his study**

**Supplementary Table S5:**
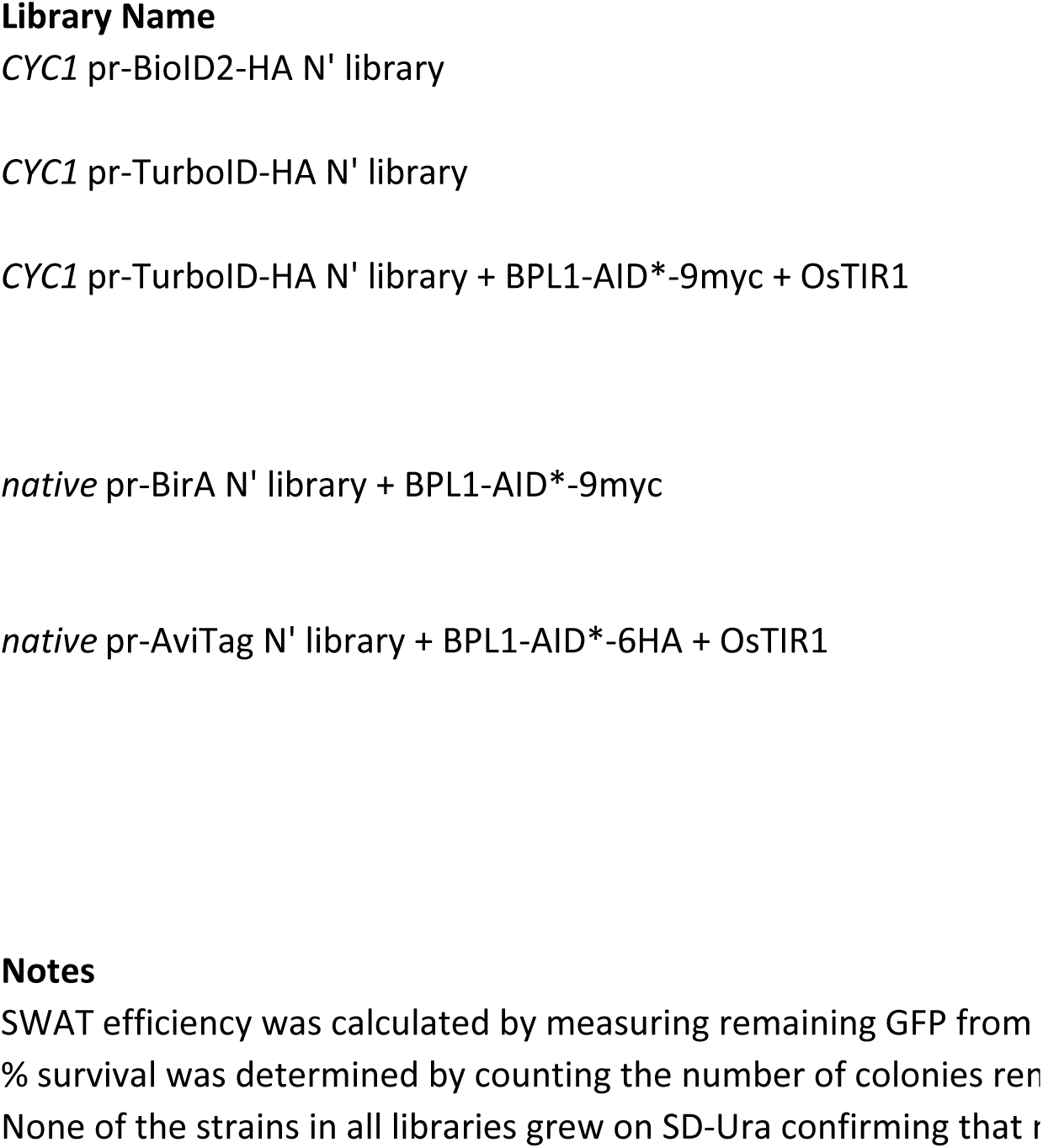
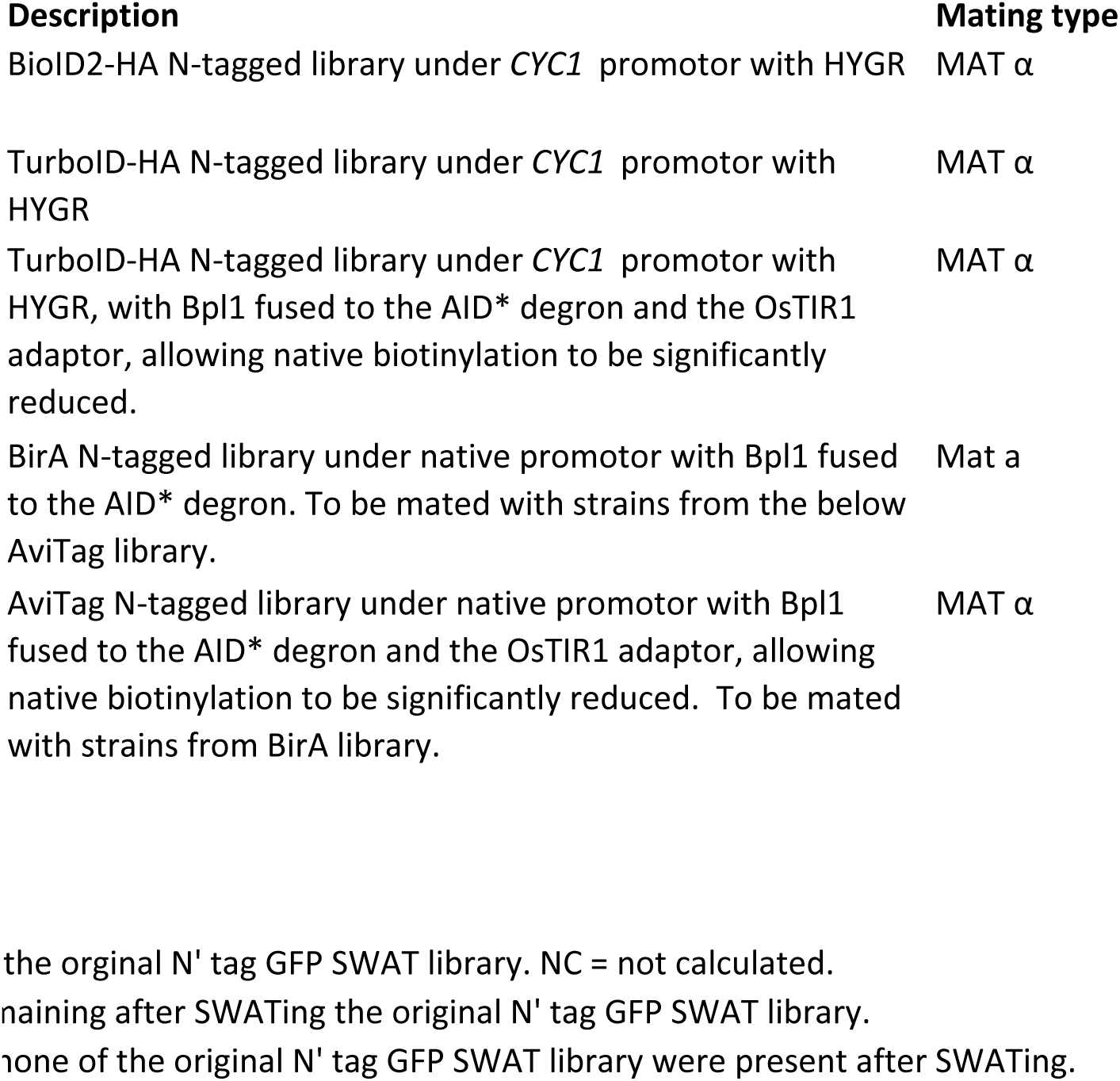
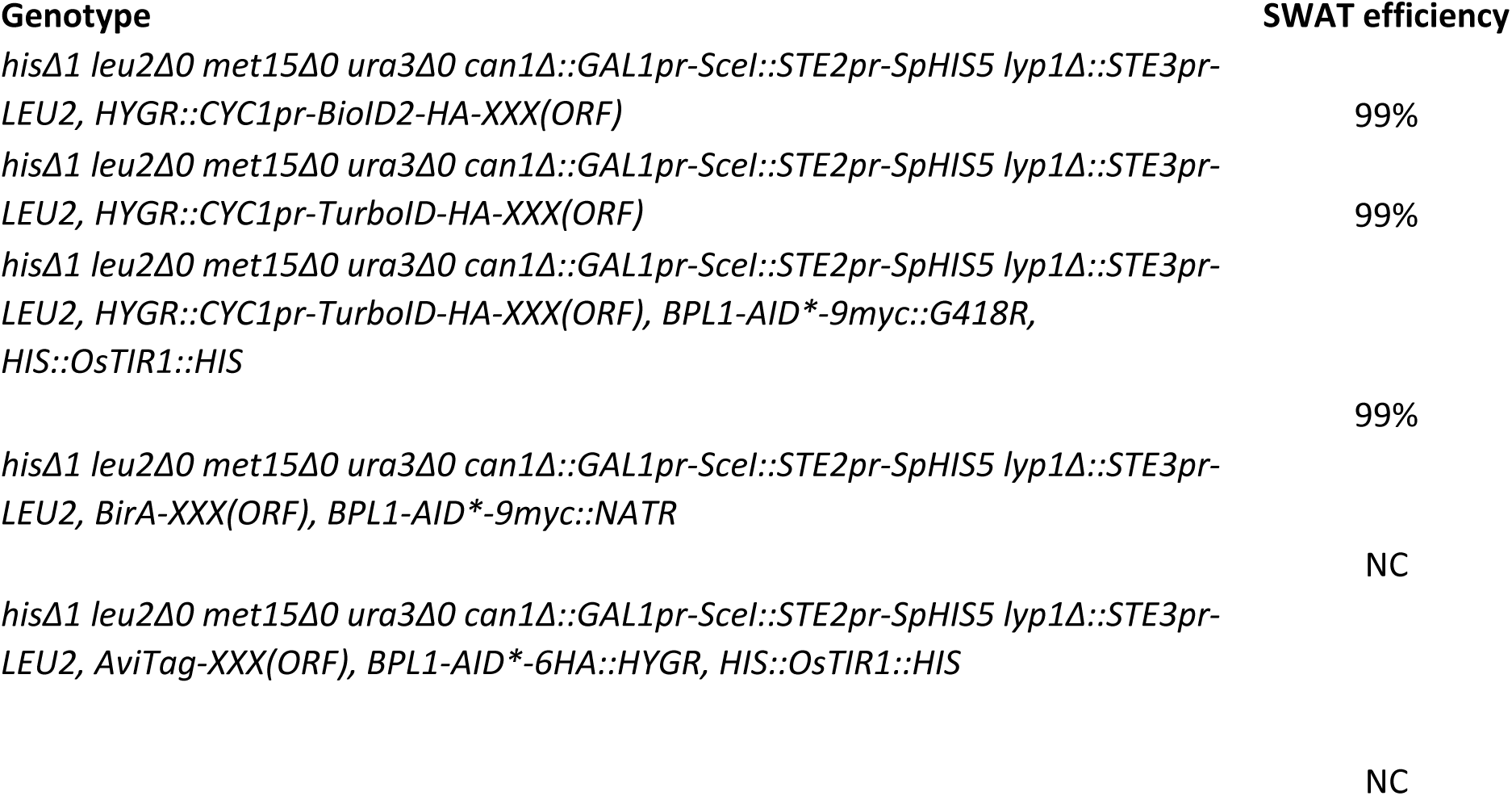
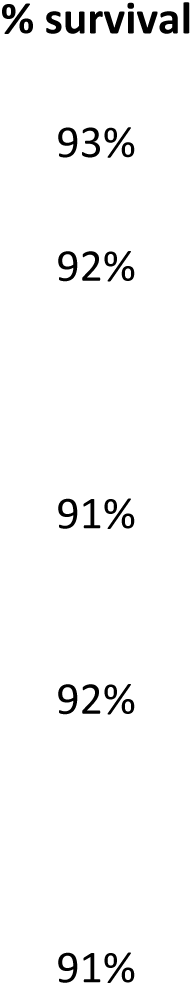
Yeast libraries generated

## Bibliography

Artan M, Barratt S, Flynn SM, Begum F, Skehel M, Nicolas A & Bono M de (2021) Interactome analysis of Caenorhabditis elegans synapses by TurboID-based proximity labeling. J Biological Chem 297: 101094

Bai L, You Q, Feng X, Kovach A & Li H (2020) Structure of the ER membrane complex, a transmembrane-domain insertase. Nature 584: 475–478

Beckett D, Kovaleva E & Schatz PJ (1999) A minimal peptide substrate in biotin holoenzyme synthetase-catalyzed biotinylation. Protein Sci 8: 921–929

Branon TC, Bosch JA, Sanchez AD, Udeshi ND, Svinkina T, Carr SA, Feldman JL, Perrimon N & Ting AY (2018) Efficient proximity labeling in living cells and organisms with TurboID. Nat Biotechnol 36: 880–887

Brewster NK, Val DL, Walker ME & Wallace JC (1994) Regulation of Pyruvate Carboxylase Isozyme (PYC1, PYC2) Gene Expression in *Saccharomyces cerevisiae* during Fermentative and Nonfermentative Growth. Archives of Biochemistry and Biophysics 311: 62–71

Chitwood PJ, Juszkiewicz S, Guna A, Shao S & Hegde RS (2018) EMC Is Required to Initiate Accurate Membrane Protein Topogenesis. Cell 175: 1507–1519.e16

Choi-Rhee E, Schulman H & Cronan JE (2004) Promiscuous protein biotinylation by Escherichia coli biotin protein ligase. Protein Science 13: 3043–3050

Christianson JC, Olzmann JA, Shaler TA, Sowa ME, Bennett EJ, Richter CM, Tyler RE, Greenblatt EJ, Harper JW & Kopito RR (2012) Defining human ERAD networks through an integrative mapping strategy. Nature Cell Biology 14: 93–105

Cronan JE (1990) Biotination of proteins in vivo. A post-translational modification to label, purify, and study proteins. J Biol Chem 265: 10327–10333

Dunham WH, Mullin M & Gingras A-C (2012) Affinity-purification coupled to mass spectrometry: Basic principles and strategies. Proteomics 12: 1576–1590

Eisenberg-Bord M, Zung N, Collado J, Drwesh L, Fenech EJ, Fadel A, Dezorella N, Bykov YS, Rapaport D, Fernandez-Busnadiego R, et al (2021) Cnm1 mediates nucleus– mitochondria contact site formation in response to phospholipid levels. J Cell Biology 220: e202104100

Fenech EJ, Ben-Dor S & Schuldiner M (2020) Double the Fun, Double the Trouble: Paralogs and Homologs Functioning in the Endoplasmic Reticulum. Annual Review of Biochemistry 89: 637–666

Gietz RD & Woods RA (2002) Transformation of yeast by lithium acetate/single-stranded carrier DNA/polyethylene glycol method. In Guide to Yeast Genetics and Molecular and Cell Biology - Part B, Fink”] [“Christine Guthrie and Gerald R. (ed) pp 87–96. Academic Press

Go CD, Knight JDR, Rajasekharan A, Rathod B, Hesketh GG, Abe KT, Youn J-Y, Samavarchi-Tehrani P, Zhang H, Zhu LY, et al (2021) A proximity-dependent biotinylation map of a human cell. Nature 595: 120–124

Guna A, Volkmar N, Christianson JC & Hegde RS (2018) The ER membrane protein complex is a transmembrane domain insertase. Science 359: 470–473

Hasslacher M, Ivessa AS, Paltauf F & Kohlwein SD (1993) Acetyl-CoA carboxylase from yeast is an essential enzyme and is regulated by factors that control phospholipid metabolism. J Biol Chem 268: 10946–10952

Hessa T, Meindl-Beinker NM, Bernsel A, Kim H, Sato Y, Lerch-Bader M, Nilsson I, White SH & Heijne G von (2007) Molecular code for transmembrane-helix recognition by the Sec61 translocon. Nature 450: 1026–1030

Hoja U, Marthol S, Hofmann J, Stegner S, Schulz R, Meier S, Greiner E & Schweizer E (2004) HFA1 Encoding an Organelle-specific Acetyl-CoA Carboxylase Controls Mitochondrial Fatty Acid Synthesis in *Saccharomyces cerevisiae*. J Biol Chem 279: 21779–21786

Ihmels J, Collins SR, Schuldiner M, Krogan NJ & Weissman JS (2007) Backup without redundancy: genetic interactions reveal the cost of duplicate gene loss. Mol Syst Biol 3: 86

Jan CH, Williams CC & Weissman JS (2014) Principles of ER cotranslational translocation revealed by proximity-specific ribosome profiling. Science 346: 1257521

Janke C, Magiera MM, Rathfelder N, Taxis C, Reber S, Maekawa H, Moreno-Borchart A, Doenges G, Schwob E, Schiebel E, et al (2004) A versatile toolbox for PCR-based tagging of yeast genes: new fluorescent proteins, more markers and promoter substitution cassettes. Yeast 21: 947–962

Jonikas MC, Collins SR, Denic V, Oh E, Quan EM, Schmid V, Weibezahn J, Schwappach B, Walter P, Weissman JS, et al (2009) Comprehensive characterization of genes required for protein folding in the endoplasmic reticulum. Science (New York, NY) 323: 1693–1697

Kim DI, Jensen SC, Noble KA, KC B, Roux KH, Motamedchaboki K & Roux KJ (2016) An improved smaller biotin ligase for BioID proximity labeling. MBoC 27: 1188–1196

Kim HS, Hoja U, Stolz J, Sauer G & Schweizer E (2004) Identification of the tRNA-binding Protein Arc1p as a Novel Target of in Vivo Biotinylation in *Saccharomyces cerevisiae*. J Biol Chem 279: 42445–42452

Kim K, Park I, Kim J, Kang M-G, Choi WG, Shin H, Kim J-S, Rhee H-W & Suh JM (2021) Dynamic tracking and identification of tissue-specific secretory proteins in the circulation of live mice. Nat Commun 12: 5204

Krahmer N, Hilger M, Kory N, Wilfling F, Stoehr G, Mann M, Farese RV & Walther TC (2013) Protein Correlation Profiles Identify Lipid Droplet Proteins with High Confidence. Mol Cell Proteomics 12: 1115–1126

Larochelle M, Bergeron D, Arcand B & Bachand F (2019) Proximity-dependent biotinylation mediated by TurboID to identify protein–protein interaction networks in yeast. J Cell Sci 132: jcs232249

Leznicki P, Schneider HO, Harvey JV, Shi WQ & High S (2021) Co-translational biogenesis of lipid droplet integral membrane proteins. J Cell Sci 135: jcs259220

Liu J, Jang JY, Pirooznia M, Liu S & Finkel T (2021) The secretome mouse provides a genetic platform to delineate tissue-specific in vivo secretion. Proc National Acad Sci 118: e2005134118

Longtine MS, McKenzie A III, Demarini DJ, Shah NG, Wach A, Brachat A, Philippsen P & Pringle JR (1998) Additional modules for versatile and economical PCR-based gene deletion and modification in *Saccharomyces cerevisiae*. Yeast 14: 953–961

Mair A, Xu S-L, Branon TC, Ting AY & Bergmann DC (2019) Proximity labeling of protein complexes and cell-type-specific organellar proteomes in Arabidopsis enabled by TurboID. Elife 8: e47864

Meurer M, Duan Y, Sass E, Kats I, Herbst K, Buchmuller BC, Dederer V, Huber F, Kirrmaier D, Stefl M, et al (2018) Genome-wide C-SWAT library for high-throughput yeast genome tagging. Nat Meth 15: 598–600

Morawska M & Ulrich HD (2013) An expanded tool kit for the auxin-inducible degron system in budding yeast. Yeast 30: 341–351

Nagaraj N, Kulak NA, Cox J, Neuhauser N, Mayr K, Hoerning O, Vorm O & Mann M (2012) System-wide Perturbation Analysis with Nearly Complete Coverage of the Yeast Proteome by Single-shot Ultra HPLC Runs on a Bench Top Orbitrap. Mol Cell Proteom Mcp 11: M111.013722

Nicastro R, Raucci S, Michel AH, Stumpe M, Osuna GMG, Jaquenoud M, Kornmann B & Virgilio CD (2021) Indole-3-acetic acid is a physiological inhibitor of TORC1 in yeast. Plos Genet 17: e1009414

Nishimura K, Fukagawa T, Takisawa H, Kakimoto T & Kanemaki M (2009) An auxin-based degron system for the rapid depletion of proteins in nonplant cells. Nat Methods 6: 917– 922

O’Keefe S, Zong G, Duah KB, Andrews LE, Shi WQ & High S (2021) An alternative pathway for membrane protein biogenesis at the endoplasmic reticulum. Commun Biology 4: 828

Opitz N, Schmitt K, Hofer-Pretz V, Neumann B, Krebber H, Braus GH & Valerius O (2017) Capturing the Asc1p/ Receptor for Activated C Kinase 1(RACK1) Microenvironment at the Head Region of the 40S Ribosome with Quantitative BioID in Yeast. Mol Cell Proteomics 16: 2199–2218

Orgil O, Matityahu A, Eng T, Guacci V, Koshland D & Onn I (2015) A Conserved Domain in the Scc3 Subunit of Cohesin Mediates the Interaction with Both Mcd1 and the Cohesin Loader Complex. Plos Genet 11: e1005036

Perez-Riverol Y, Bai J, Bandla C, García-Seisdedos D, Hewapathirana S, Kamatchinathan S, Kundu DJ, Prakash A, Frericks-Zipper A, Eisenacher M, et al (2021) The PRIDE database resources in 2022: a hub for mass spectrometry-based proteomics evidences. Nucleic Acids Res 50: D543–D552

Pirner HM & Stolz J (2006) Biotin Sensing in *Saccharomyces cerevisiae* is Mediated by a Conserved DNA Element and Requires the Activity of Biotin-Protein Ligase. J Biol Chem 281: 12381–12389

Roux KJ, Kim DI, Raida M & Burke B (2012) A promiscuous biotin ligase fusion protein identifies proximal and interacting proteins in mammalian cells. J Cell Biology 196: 801– 810

Sanchez AD, Branon TC, Cote LE, Papagiannakis A, Liang X, Pickett MA, Shen K, Jacobs-Wagner C, Ting AY & Feldman JL (2021) Proximity labeling reveals non-centrosomal microtubule-organizing center components required for microtubule growth and localization. Curr Biol 31: 3586–3600.e11

Schindelin J, Arganda-Carreras I, Frise E, Kaynig V, Longair M, Pietzsch T, Preibisch S, Rueden C, Saalfeld S, Schmid B, et al (2012) Fiji: an open-source platform for biological-image analysis. Nat Methods 9: 676–682

Shurtleff MJ, Itzhak DN, Hussmann JA, Oakdale NTS, Costa EA, Jonikas M, Weibezahn J, Popova KD, Jan CH, Sinitcyn P, et al (2018) The ER membrane protein complex interacts cotranslationally to enable biogenesis of multipass membrane proteins. Elife 7: e37018

Singh AP, Salvatori R, Aftab W, Aufschnaiter A, Carlström A, Forne I, Imhof A & Ott M (2020) Molecular Connectivity of Mitochondrial Gene Expression and OXPHOS Biogenesis. Mol Cell 79: 1051–1065.e10

Sumrada RA & Cooper TG (1982) Urea carboxylase and allophanate hydrolase are components of a multifunctional protein in yeast. J Biol Chem 257: 9119–9127

Tang X, Snowball JM, Xu Y, Na C-L, Weaver TE, Clair G, Kyle JE, Zink EM, Ansong C, Wei W, et al (2017) EMC3 coordinates surfactant protein and lipid homeostasis required for respiration. J Clin Invest 127: 4314–4325

Tian S, Wu Q, Zhou B, Choi MY, Ding B, Yang W & Dong M (2019) Proteomic Analysis Identifies Membrane Proteins Dependent on the ER Membrane Protein Complex. Cell Reports 28: 2517–2526.e5

Tong AHY & Boone C (2007) 16 High-Throughput Strain Construction and Systematic Synthetic Lethal Screening in *Saccharomyces cerevisiae*. In Yeast Gene Analysis, Stark”] [“Ian Stansfield and Michael JR (ed) pp 369–707. Academic Press

Uçkun E, Wolfstetter G, Anthonydhason V, Sukumar SK, Umapathy G, Molander L, Fuchs J & Palmer RH (2021) In vivo Profiling of the Alk Proximitome in the Developing *Drosophila* Brain. J Mol Biol 433: 167282

Uezu A, Kanak DJ, Bradshaw TWA, Soderblom EJ, Catavero CM, Burette AC, Weinberg RJ & Soderling SH (2016) Identification of an elaborate complex mediating postsynaptic inhibition. Science 353: 1123–1129

Volkmar N, Thezenas M-L, Louie SM, Juszkiewicz S, Nomura DK, Hegde RS, Kessler BM & Christianson JC (2019) The ER membrane protein complex promotes biogenesis of sterol-related enzymes maintaining cholesterol homeostasis. J Cell Sci 132: jcs223453

Weill U, Cohen N, Fadel A, Ben-Dor S & Schuldiner M (2019) Protein Topology Prediction Algorithms Systematically Investigated in the Yeast *Saccharomyces cerevisiae*. BioEssays 41: 1800252

Weill U, Yofe I, Sass E, Stynen B, Davidi D, Natarajan J, Ben-Menachem R, Avihou Z, Goldman O, Harpaz N, et al (2018) Genome-wide SWAp-Tag yeast libraries for proteome exploration. Nat Meth: 1–13

Yofe I & Schuldiner M (2014) Primers-4-Yeast: a comprehensive web tool for planning primers for *Saccharomyces cerevisiae*. Yeast 31: 77–80

Yofe I, Weill U, Meurer M, Chuartzman S, Zalckvar E, Goldman O, Ben-Dor S, Schütze C, Wiedemann N, Knop M, et al (2016) One library to make them all: streamlining the creation of yeast libraries via a SWAp-Tag strategy. Nat Meth 13: 371–378

Zhang Y, Song G, Lal NK, Nagalakshmi U, Li Y, Zheng W, Huang P, Branon TC, Ting AY, Walley JW, et al (2019) TurboID-based proximity labeling reveals that UBR7 is a regulator of N NLR immune receptor-mediated immunity. Nat Commun 10: 3252

